# Prenatal Cannabidiol and Δ9-Tetrahydrocannabinol exposure lead to sex-specific disruptions in risk assessment and behavioral switching via divergent rewiring of the adult mPFC

**DOI:** 10.64898/2026.06.27.734813

**Authors:** Alba Cáceres-Rodríguez, Daniela Iezzi, Olivier Lassalle, Anna E. Dudek, Shuai Wang, Pascale Chavis, Olivier J. Manzoni

## Abstract

**Background:** Prenatal cannabidiol (CBD) consumption is increasing, driven by a perception of safety relative to Δ9-tetrahydrocannabinol (THC). However, the neurodevelopmental risks of gestational CBD remain largely uncharacterized.

**Methods:** Using a sex-disaggregated framework, we investigated adult (P100–140) mouse offspring following in utero exposure (GD5-18; 3 mg/kg) to THC or CBD. Behavioral strategies were evaluated through risk-assessment and repetitive behavior tasks, coupled with targeted electrophysiological mapping of medial prefrontal cortex Layer 5 neurons, the primary hub for approach-avoidance arbitration.

**Results:** We found that increased repetitive behavior was a universal feature of prenatal cannabinoids exposure. Alterations in risk appraisal emerged uniquely in CBD-exposed females and appeared dissociated from classical anxiety metrics. At the circuit level, THC and CBD were linked to an absence of endocannabinoid long-term depression (eCB-LTD). In males, CBD exposure coincided with a bidirectional plasticity collapse characterized by functional saturation, elevated AMPA/NMDA ratios, and slowed NMDAR activation kinetics. This ceiling effect may represent a top-down constraint on the prefrontal output circuit, potentially limiting the synaptic flexibility typically associated with adaptive behavioral transitions. In contrast, females exhibited compound-specific reorganizations of E/I balance. CBD-exposed females displayed a scaled-up architecture that preserved net E/I balance, whereas THC was associated with a pro-excitatory phenotype through the collapse of inhibitory control.

**Conclusions:** Despite a shared loss of eCB-LTD, distinct synaptic remodeling might underlie divergent alterations in risk assessment and behavioral flexibility. This sex-specific circuit rewiring provides a neurobiological framework for the long-term behavioral risks associated with gestational cannabinoid exposure.

## Introduction

Amid expanding legalization, gestational cannabis use has surged, with cannabidiol (CBD) widely framed as a natural alternative to THC (1). Yet, its adoption has outpaced neurobiological evidence. Unlike the documented risks of prenatal THC, the long-term impact of gestational CBD on adult brain function remains poorly understood, presenting a pressing public health concern. As a master regulator of mammalian neurodevelopment, the endocannabinoid system orchestrates neuronal migration, axonal pathfinding, and circuit refinement (2–4). Over the past decade, prenatal cannabinoid exposure (PCE) -primarily via THC or synthetic agonists-has been shown to induce persistent, male-biased social deficits and altered socio-affective trajectories (5–7). Mechanistically, we and others have linked these impairments to the miswiring of critical developmental hubs, including a hyperdopaminergic phenotype in the ventral tegmental area (8), the permanent loss of hippocampal cholecystokinin-containing interneurons (7), and the ablation of endocannabinoid-mediated long-term depression (eCB-LTD) in the medial prefrontal cortex (5,9,10).

Unlike documented perinatal THC phenotypes, the lasting neurobehavioral impact of gestational CBD remains largely uncharacterized. Recent data challenge CBD’s assumed neutrality, showing fetal exposure alters thermal sensitivity (11), disrupts early communication (12), and selectively impairs adult cognition (13). We recently established that gestational CBD permanently remodels the adult insular cortex, yielding progressive anxiety-like phenotypes (14,15). Yet, whether prenatal cannabinoid exposure fundamentally impairs the risk-evaluation strategies and motor-cognitive transitions required for complex environmental adaptation remains unknown.

Risk appraisal and motor flexibility demand top-down sensory integration to modulate reflexive outputs (16–18). Whether gestational CBD or THC disrupts approach-avoidance arbitration or behavioral state transitions remains unresolved. We hypothesized that these vulnerabilities manifest in specialized ethological readouts: Stretched Attend Posture (SAP) in the elevated plus maze (EPM) to quantify active risk assessment, and repetitive motor persistence in the marble burying (MB) task to probe behavioral switching.

Because mPFC layer-5 pyramidal output neurons gate risk-assessment and behavioral switching, we targeted this circuit to isolate the cellular substrates driving these behavioral shifts. Providing a sex-disaggregated comparison, we report that prenatal THC and CBD both induce profound synaptic rigidity, characterized by a bidirectional collapse of plasticity. This circuit-specific failure proposes a mechanistic basis for disrupted risk appraisal and behavioral switching, rather than a generalized adaptation deficit. Collectively, our data demonstrate that CBD replicates the synaptic and behavioral vulnerabilities long attributed to THC, establishing a common signature of dysfunction that strengthens the rationale against gestational CBD use.

## Materials and methods

### Animals

Animal protocols were approved by the French Ethical Committee (APAFIS #49376 and #49376-2024051414491391) in accordance with European Council Directive (86/609/EEC) and NIH guidelines. C57BL/6J mice (Charles River) were maintained under standard colony conditions. At gestational day 0 (GD0, identified by vaginal plug), dams were individually housed. From GD5-GD18, dams received daily subcutaneous injections (4 mL/kg) of vehicle (1:1:18 Cremophor EL:ethanol:saline), CBD (3 mg/kg), or THC (3 mg/kg)(NIDA Drug Supply Program) (12,14,15). Offspring were weaned at PND21 and housed by sex until adulthood (PND90–140). To eliminate litter bias without culling, groups comprised full offspring from 3-4 litters.

### Behavior

Adult anxiety and repetitive behaviors were assessed via EPM (including spatial profiling of posture) (15,19–21) and MB (22) using EthoVision (Noldus). Normal data (D’Agostino–Pearson; Shapiro–Wilk) were analyzed in GraphPad Prism 11 via two-way ANOVA with Šídák’s post-hoc test; the ROUT method removed outliers. Full statistics are in figure legends and tables.

### Physiology

#### Slice Preparation

Adult mice (PND 100–140) were anesthetized with isoflurane and sacrificed. Coronal PFC slices (300 μm; Campden Instruments vibratome) were prepared in ice-cold sucrose cutting solution (in mM: 87 NaCl, 75 sucrose, 25 glucose, 2.5 KCl, 4 MgCl₂, 0.5 CaCl₂, 23 NaHCO₃, 1.25 NaH₂PO₄) (21–23). Slices recovered for 30 min at 32°C in oxygenated (95% O₂ / 5% CO₂) low-calcium aCSF (in mM: 130 NaCl, 11 glucose, 2.5 KCl, 2.4 MgCl₂, 1.2 CaCl₂, 23 NaHCO₃, 1.2 NaH₂PO₄) and were kept at room temperature until recording (23–25).

#### Electrophysiology

Whole-cell patch-clamp recordings were obtained from prelimbic PFC layer 5/6 pyramidal neurons as previously described (23–28). Slices were continuously perfused (2 mL/min, 32°C) with low-calcium solution containing 10 μM gabazine. Data were recorded using an Axopatch-200B amplifier, digitized via DigiData 1440A, and analyzed with Clampex 10.7 (Molecular Devices. Patch pipettes (2–3 MΩ; access resistance < 25 MΩ) were filled with an intracellular solution containing (in mM): 1145 K+ gluconate, 3 NaCl, 1 MgCl2, 1 EGTA, 0.3 CaCl2, 2 Na2+ ATP, 0.3 Na+ GTP, and 0.2 cAMP, buffered with 10 HEPES (pH 7.25, 290–300 mOsm). Neurons were held at −70 mV. Immediately following breakthrough, current-voltage (I-V) relationships were established using a series of hyperpolarizing-to-depolarizing current steps.

##### Postsynaptic Spontaneous Activity and Evoked Plasticity

Spontaneous AMPA, NMDA, and GABA currents were pharmacologically isolated (14,27). AMPA-sEPSCs: Recorded at −70 mV using the K-gluconate internal solution with 10 µM gabazine 10 µM, SR 95531 hydrobromide; Tocris). GABA-sIPSCs: Recorded at −70 mV in the presence of 20 µM CNQX and 50 µM APV (Tocris), using a high-chloride internal solution (in mM: 140 KCl, 1.6 MgCl₂, 2.5 MgATP, 0.5 NaGTP, 2 EGTA, 10 HEPES; pH 7.25–7.3, 280–300 mOsm). NMDA-sEPSCs: Recorded at +30 mV in the presence of 20 µM NBQX (Tocris), using a cesium-methanesulfonate internal solution (in mM: 143 Cs-MeSO₃, 5 NaCl, 1 MgCl₂, 1 EGTA, 0.3 CaCl₂, 10 HEPES, 2 Na₂ATP, 0.3 NaGTP, 0.2 cAMP; pH 7.3, 290 mOsm).

##### AMPA/NMDA Ratio

Evoked EPSCs were recorded from layer 5 pyramidal neurons at +30 mV (Cs-methanesulfonate internal) via layer 2/3 electrical stimulation. The NMDA component was isolated by bath application of 20 μM NBQX, and the AMPA component was derived via digital subtraction (27).

##### Extracellular Recordings

Layer 5 Field excitatory postsynaptic potentials (fEPSP) were evoked by layer 2/3 stimulation (0.1 Hz, 60% maximum intensity); glutamatergic transmission was confirmed using 10 µM CNQX. LTP was induced via theta-burst stimulation (TBS: 5 trains of 4 pulses at 100 Hz, 200-ms interval, repeated 4 times every 10 s) (27), and eCB-LTD via 10-Hz stimulation for 10 min (24).

### Data Analysis and Statistics

Data were analyzed using Clampfit 11.3, AxoGraph 1.7.6, and GraphPad Prism 10.4.1. Resting membrane potential (RMP) was measured at breakthrough. Membrane capacitance (Cm) was calculated by integrating capacitive currents (−2 mV pulse), and input resistance from the I-V relationship near RMP. Intrinsic excitability curves were generated via depolarizing current steps, with rheobase and threshold determined using 10-pA increments. Spontaneous currents were template-detected in AxoGraph (threshold: 3.5×baseline noise SD; minimum 7 pA) with specific rise/decay constants (ms): AMPA (0.5/3), GABA (0.2/10), and NMDA (3/10). Total charge transfer was integrated over 6 min (15,27). The AMPA/NMDA ratio was calculated at +30 mV following digital subtraction of the NMDA component. Field plasticity was quantified 30–40 min post-induction relative to baseline. Following normality (D’Agostino-Pearson; Shapiro-Wilk) and outlier (ROUT) screening, data were evaluated by two-way ANOVA with Šídák’s post-hoc test and visualized as violin plots with individual data points.

#### Synaptic Distribution and E/I Balance Analysis

Statistical analyses were performed using R (v4.2.2) and GraphPad Prism (v11). Cumulative distributions of synaptic charge transfer were compared via Kolmogorov-Smirnov tests. To evaluate net functional balance, an E/I index (E/[E+I]) was computed. Because E and I currents were recorded from independent neuronal populations, a nonparametric bootstrap (10,000 iterations with replacement) was executed in R to generate stable index distributions. Group differences (ΔE/I index) were quantified by subtracting these bootstrap distributions (e.g., Sham minus cannabinoid groups). Significance was established if more than 95% of the ΔE/I distribution fell on one side of zero (p < 0.05). Full statistical parameters are tabulated; figures isolate key cross-sex control and within-sex treatment comparisons.

## Results

Daily subcutaneous cannabinoid exposure (GD5–18) did not alter maternal weight gain, gestational length, litter size, or sex ratios, confirming normal pregnancy progression (Table 1).

**Table 1:** Follow up of gestational parameters and litter features. Statistical significance defined as p-value < 0.05.

| Gestational parameters | Treatment | Median | Max | Min | N | Kruskal-Wallis test (p-value) |
| --- | --- | --- | --- | --- | --- | --- |
| Gestational weight gain (grams) | SHAM | 16.40 | 18.70 | 12.70 | 18 | 0.4491 |
|  | CBD | 16.95 | 20.70 | 13.10 | 24 |  |
|  | THC | 16.80 | 19.80 | 11.50 | 9 |  |
| Duration of gestation (days) | SHAM | 19.00 | 20.00 | 18.00 | 18 | 0.7293 |
|  | CBD | 19.00 | 20.00 | 17.00 | 24 |  |
|  | THC | 19.00 | 19.00 | 18.00 | 9 |  |
| Litter size (#) | SHAM | 7.00 | 10.00 | 3.00 | 18 | 0.0813 |
|  | CBD | 8.00 | 10.00 | 1.00 | 24 |  |
|  | THC | 8.00 | 10.00 | 5.00 | 9 |  |
| Ratio males/females | SHAM | 1.33 | 5.00 | 0.33 | 18 | 0.1697 |
|  | CBD | 1.63 | 9.00 | 0.14 | 25 |  |
|  | THC | 0.67 | 4.00 | 0.17 | 9 |  |

### Prenatal cannabinoids spare classical adult anxiety indices despite adolescent vulnerability

Although epidemiological and preclinical data suggest that in utero cannabis exposure increases the risk for affective disorders (29–32), the stability and divergence of specific prenatal CBD and THC phenotypes remain poorly defined. Extending our finding of a female-specific anxiogenic phenotype in CBD-exposed adolescents (12,15), we mapped adult EPM profiles. Canonical anxiety metrics revealed no significant sex or treatment effects (Table 2, Supplemental Fig. 1); however, prenatal THC selectively induced hyper-locomotion in adult females. Crucially, the adolescent anxiety phenotype in CBD females resolved in adulthood, suggesting a developmental normalization of baseline approach-avoidance exploration consistent with prior reports of stable open-arm exploration following prenatal THC (5).

**Table 2:** Coventional EPM analysis. Statistical significance defined as p-value < 0.05.

| Measure | Treatment | Median | Max | Min | N | Two-way ANOVA Main effects and interactions | Multiple comparison (Šidák's post hoc test) |
| --- | --- | --- | --- | --- | --- | --- | --- |
| Distance moved (cm) | SHAM FEMALE | 957.618 | 1591.55 | 503.066 | 13 | **Interaction:** F (2, 74) = 0,3939; P=0,6758<br>**Sex:** F (1, 74) = 1,398; P=0,2409<br>**Treatment** F (2, 74) = 4,419; P=0,0154 | FEMALE<br>"Sham vs. CBD" P=0,2099<br>"Sham vs. THC" P=0,0250<br>"CBD vs. THC" P=0,6904<br>MALE<br>"Sham vs. CBD" P=0,8485<br>"Sham vs. THC" P=0,2752<br>"CBD vs. THC" P=0,6260 |
|  | CBD FEMALE | 1150.015 | 1723.54 | 750.624 | 12 |  |  |
|  | THC FEMALE | 1282.48 | 1661.58 | 899.255 | 16 |  |  |
|  | SHAM MALE | 1075.535 | 2020.9 | 810.543 | 11 |  |  |
|  | CBD MALE | 1282.85 | 1758.49 | 584.972 | 10 |  |  |
|  | THC MALE | 1276.36 | 1705.87 | 1012.67 | 16 |  |  |
| Preference for Open Arms (%) | SHAM FEMALE | 17.39257 | 58.8596 | 0.383216 | 13 | **Interaction:** F (2, 72) = 0,3889; P=0,6792<br>**Sex:** F (1, 72) = 0,1030; P=0,7492<br>**Treatment** F (2, 72) = 2,464; P=0,0922 | FEMALE<br>"Sham vs. CBD" P=0,9909<br>"Sham vs. THC" P=0,4119<br>"CBD vs. THC" P=0,5080<br>MALE<br>"Sham vs. CBD" P=0,3921<br>"Sham vs. THC" P=0,1629<br>"CBD vs. THC" P=0,9285 |
|  | CBD FEMALE | 25.64002 | 49.27898 | 5.156046 | 12 |  |  |
|  | THC FEMALE | 28.300745 | 93.13346 | 7.419347 | 16 |  |  |
|  | SHAM MALE | 17.72665 | 42.86245 | 3.019399 | 11 |  |  |
|  | CBD MALE | 33.124375 | 55.86683 | 7.895273 | 10 |  |  |
|  | THC MALE | 33.847205 | 71.15472 | 3.166812 | 16 |  |  |
| Closed Arms + Centre (%) | SHAM FEMALE | 83.4482 | 99.41843 | 62.6866 | 13 | **Interaction:** F (2, 72) = 0,2805; P=0,7563<br>**Sex:** F (1, 72) = 0,2130; P=0,9357<br>**Treatment** F (2, 72) = 2,798; P=0,0676 | FEMALE<br>"Sham vs. CBD" P=0,9243<br>"Sham vs. THC" P=0,3009<br>"CBD vs. THC" P=0,5406<br>MALE<br>"Sham vs. CBD" P=0,3705<br>"Sham vs. THC" P=0,1621<br>"CBD vs. THC" P=0,9430 |
|  | CBD FEMALE | 79.02395 | 93.5023 | 66.329 | 12 |  |  |
|  | THC FEMALE | 76.99995 | 92.7134 | 51.2264 | 16 |  |  |
|  | SHAM MALE | 84.0974 | 96.83627 | 66.5836 | 11 |  |  |
|  | CBD MALE | 74.4891 | 91.90925 | 63.9245 | 10 |  |  |
|  | THC MALE | 73.7754 | 95.84524 | 56.1791 | 16 |  |  |
| Open Arms (%) | SHAM FEMALE | 15.09575 | 37.0318 | 0.387722 | 13 | **Interaction:** F (2, 75) = 0,8071; P=0,4500<br>**Sex:** F (1, 75) = 0,006545; P=0,6458<br>**Treatment** F (2, 75) = 2,087; P=0,1312 | FEMALE<br>"Sham vs. CBD" P=0,9997<br>"Sham vs. THC" P=0,7523<br>"CBD vs. THC" P=0,7379<br>MALE<br>"Sham vs. CBD" P=0,9997<br>"Sham vs. THC" P=0,7523<br>"CBD vs. THC" P=0,7379 |
|  | CBD FEMALE | 20.3919 | 33.0084 | 4.90141 | 12 |  |  |
|  | THC FEMALE | 22.04415 | 48.213 | 0.381558 | 16 |  |  |
|  | SHAM MALE | 14.3997 | 29.9907 | 0 | 11 |  |  |
|  | CBD MALE | 24.776 | 35.8404 | 7.31569 | 10 |  |  |
|  | THC MALE | 25.2479 | 42.2859 | 3.07094 | 16 |  |  |

### Prenatal CBD, but not THC, induces enduring female-specific increases in risk-assessment postures

To determine if prenatal cannabinoid exposure alters active exploration, we performed an ethological analysis of postural states during the EPM (Figure 1, Table 3). While a neutral stance was the predominant posture across all groups, a baseline sexual dimorphism emerged: males consistently expressed neutral postures more frequently than treatment-matched females. Within females, THC exposure significantly increased neutral stances compared to Sham controls. Crucially, tracking SAP, a validated index of risk assessment, unveiled a selective vulnerability in the CBD lineage. CBD-exposed females engaged in SAP significantly longer than both Sham females and CBD males, signaling heightened hesitation. This strategy shift was strictly compound-specific, absent in all THC groups, and accompanied by a reciprocal reduction in contracted postures.

**Figure 1.**
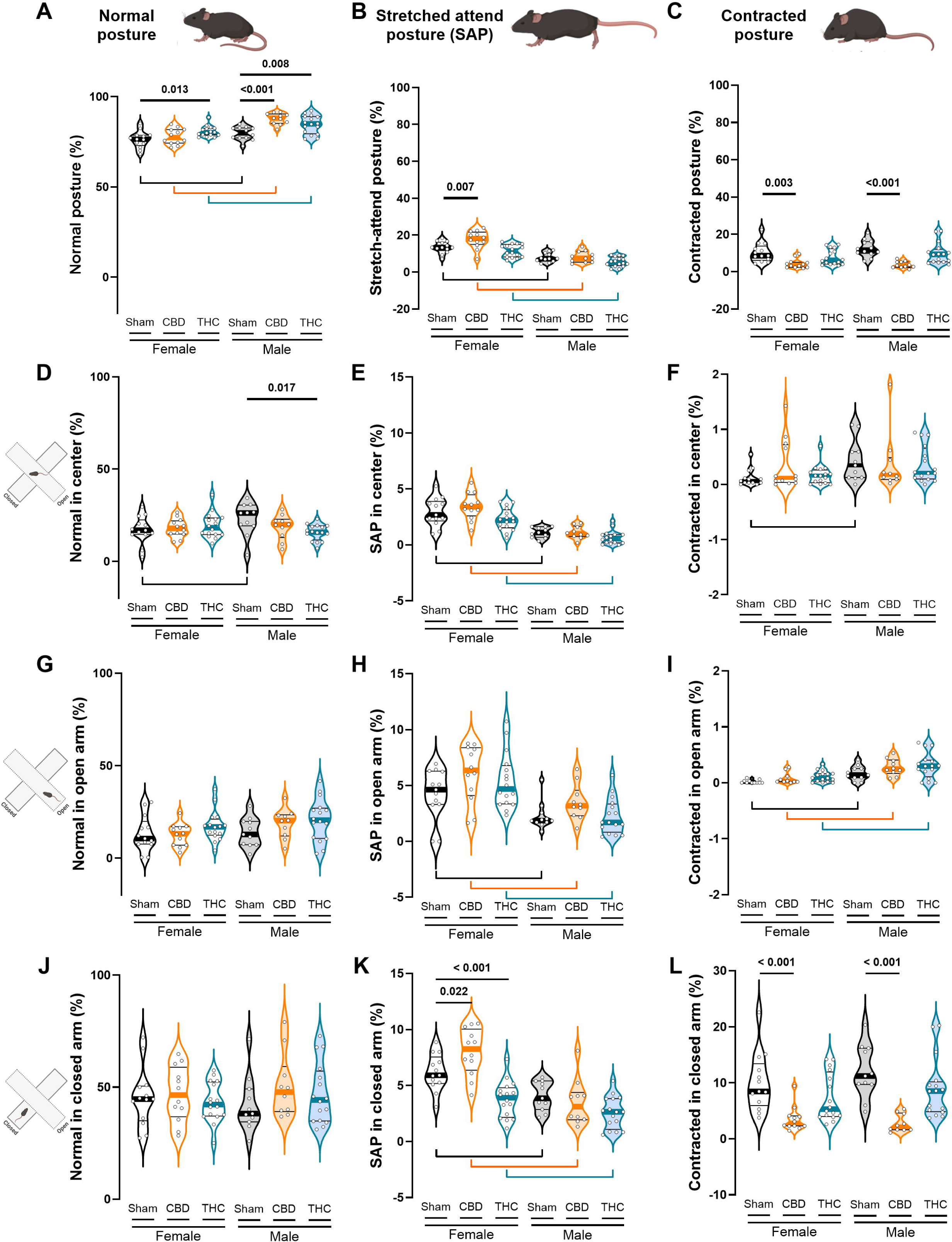
Prenatal exposure to CBD, but not THC, selectively enhances risk assessment behavior in adult female offspring. Postural analysis during the EPM task categorized behavior into three distinct states: normal posture (A), stretched-attend posture (SAP; B), and contracted posture (C). (A) Female offspring exposed to prenatal THC exhibited a higher prevalence of normal posture. (B) Analysis revealed that CBD-exposed females spent significantly more time in SAP compared to both Sham females and CBD-exposed males — an effect absent in THC-exposed groups. Female offspring in general display higher incidence of SAP than males, this is preserved across within treatment sex comparisons (C) CBD-exposed animals also spent less time in contracted posture, consistent with an increased expression of SAP-related risk assessment behavior. Statistical comparisons were performed using two-way ANOVA followed by Šídák’s multiple comparisons test. (D–L) Quantitative analysis of postural states, normal, contracted, and stretched-attend posture (SAP), across the three zones of the EPM revealed that, regardless of sex or prenatal treatment, both Sham and CBD-exposed offspring spent significantly more time in normal posture within the closed arms compared to the open and center arms. Notably, CBD-exposed progeny, particularly females, exhibited elevated SAP rates specifically from the closed arms prior to center entry. Data are presented as violin plots (median, and 25th–75th percentiles), with each point representing an individual animal. Statistical analysis was performed using two-way ANOVA followed by Šídák’s multiple comparisons test. *P values < 0.05 are indicated in the graph. Group sizes were as follows: Sham female (N = 13), CBD female (N = 12), THC female (N = 16), Sham male (N = 11), CBD male (N = 10), THC male (N = 16).

**Table 3:** Postural analysis during EPM. Statistical significance defined as p-value < 0.05.

| Measure | Treatment | Median | Max | Min | N | Two-way ANOVA<br>Main effects and<br>interactions | Multiple comparison<br>(Šidák's post hoc<br>test) |
| --- | --- | --- | --- | --- | --- | --- | --- |
| Normal<br>posture (%) | SHAM FEMALE | 76.44 | 85.2<br>3 | 68.15 | 13 | **Interaction:** F (2, 74) =<br>4,760; P=0,0114<br>**Sex:** F (1, 74) = 38,20;<br>P<0,0001<br>**Treatment** F (2, 74) =<br>11,64; P<0,0001 | **FEMALE**<br>"Sham vs. CBD" P=0,5584<br>"Sham vs. THC" P=0,0125<br>"CBD vs. THC" P=0,1842<br>**MALE**<br>"Sham vs. CBD" P<0,0001<br>"Sham vs. THC" P=0,0078<br>"CBD vs. THC" P=0,1059<br>**FEMALE vs. MALE**<br>"Sham" P=0,0386<br>"CBD" P<0,0001<br>"THC" P=0,0103 |
|  | CBD FEMALE | 77.40 | 84.7<br>7 | 71.86 | 12 |  |  |
|  | THC FEMALE | 80.09 | 88.5<br>7 | 77.29 | 16 |  |  |
|  | SHAM MALE | 80.02 | 84.8<br>9 | 72.60 | 11 |  |  |
|  | CBD MALE | 88.15 | 91.5<br>4 | 81.81 | 10 |  |  |
|  | THC MALE | 84.97 | 92.2<br>1 | 76.58 | 16 |  |  |
| Strech attend<br>posture (%) | SHAM FEMALE | 13.29 | 18.4<br>6 | 7.14 | 13 | **Interaction:** F (2, 74) =<br>2,479; P=0,0908<br>**Sex:** F (1, 74) = 87,79;<br>P<0,0001<br>**Treatment** F (2, 74) =<br>11,82; P<0,0001 | **FEMALE**<br>"Sham vs. CBD" P=0,0064<br>"Sham vs. THC" P=0,2608<br>"CBD vs. THC" P<0,0001<br>**MALE**<br>"Sham vs. CBD" P=0,9611<br>"Sham vs. THC" P=0,1769<br>"CBD vs. THC" P=0,1098<br>**FEMALE vs. MALE**<br>"Sham" P<0,0001<br>"CBD" P<0,0001<br>"THC" P<0,0001 |
|  | CBD FEMALE | 18.06 | 23.9<br>7 | 7.08 | 12 |  |  |
|  | THC FEMALE | 11.40 | 16.4<br>3 | 6.21 | 16 |  |  |
|  | SHAM MALE | 7.27 | 11.8<br>8 | 4.52 | 11 |  |  |
|  | CBD MALE | 7.41 | 13.8<br>0 | 4.33 | 10 |  |  |
|  | THC MALE | 5.31 | 9.25 | 1.60 | 16 |  |  |
| Contracted<br>posture (%) | SHAM FEMALE | 8.54 | 22.5<br>9 | 4.28 | 13 | **Interaction:** F (2, 74) =<br>0,7749; P=0,4645<br>**Sex:** F (1, 74) = 1,642;<br>P=0,2040<br>**Treatment** F (2, 74) =<br>17,64; P<0,0001 | **FEMALE**<br>"Sham vs. CBD" P=0,0026<br>"Sham vs. THC" P=0,2678<br>"CBD vs. THC" P=0,0982<br>**MALE**<br>"Sham vs. CBD" P<0,0001<br>"Sham vs. THC" P=0,2682<br>"CBD vs. THC" P=0,0017<br>**FEMALE vs. MALE**<br>"Sham" P=0,2143<br>"CBD" P=0,7723<br>"THC" P=0,1713 |
|  | CBD FEMALE | 3.55 | 9.61 | 1.79 | 12 |  |  |
|  | THC FEMALE | 6.12 | 14.5<br>4 | 3.29 | 16 |  |  |
|  | SHAM MALE | 11.58 | 20.9<br>0 | 5.67 | 11 |  |  |
|  | CBD MALE | 3.29 | 6.93 | 1.21 | 10 |  |  |
|  | THC MALE | 9.26 | 21.8<br>2 | 3.84 | 16 |  |  |

Spatial mapping revealed that this postural phenotype was not uniform but zone-restricted (Figure 1, Table 4). While all cohorts spent the majority of the time in neutral-posture within the closed arms, the elevated SAP in CBD females was localized entirely within the closed arms, displaying increased exploratory behavior or hesitation immediately prior to entering the center/open arm zone. This spatial bias demonstrates that prenatal CBD selectively disrupts the preparatory phase of exploration, promoting hyper-vigilance within relative safety before approaching a zone of uncertainty.

**Table 4:** Postural analysis during EPM per area of the maze. Statistical significance defined as p-value < 0.05.

| Measure | Area | Treatment | Median | Max | Min | N | Two-way ANOVA<br>Main effects and<br>interactions | Multiple comparison<br>(Šidák's post hoc<br>test) |
| --- | --- | --- | --- | --- | --- | --- | --- | --- |
| Normal<br>posture (%) | Center | SHAM FEMALE | 16.9 | 29.5 | 2.4 | 13 | **Interaction:** F (2, 72) = 3,582; P=0,0329<br>**Sex:** F (1, 72) = 0,3666; P=0,5468<br>**Treatment** F (2, 72) = 0,9996; P=0,3731 | **FEMALE**<br>"Sham vs. CBD" P=0,9334<br>"Sham vs. THC" P=0,6266<br>"CBD vs. THC" P=0,8553<br>**MALE**<br>"Sham vs. CBD" P=0,2775<br>"Sham vs. THC" P=0,0170<br>"CBD vs. THC" P=0,5388<br>**FEMALE vs. MALE**<br>"Sham" P=0,0324<br>"CBD" P=0,8552<br>"THC" P=0,1212 |
|  |  | CBD FEMALE | 17.9 | 26.5 | 10.5 | 12 |  |  |
|  |  | THC FEMALE | 18.3 | 36.1 | 9.7 | 16 |  |  |
|  |  | SHAM MALE | 26.2 | 34.3 | 3.5 | 11 |  |  |
|  |  | CBD MALE | 19.9 | 28.2 | 6.6 | 10 |  |  |
|  |  | THC MALE | 15.8 | 22.9 | 7.1 | 16 |  |  |
|  | Open<br>arms | SHAM FEMALE | 10.5 | 30.3 | 0.4 | 13 | **Interaction:** F (2, 72) = 0,5525; P=0,5779<br>**Sex:** F (1, 72) = 2,863; P=0,0949<br>**Treatment** F (2, 72) = 2,717; P=0,0729 | x |
|  |  | CBD FEMALE | 13.2 | 24.3 | 2.9 | 12 |  |  |
|  |  | THC FEMALE | 16.8 | 37.2 | 4.1 | 16 |  |  |
|  |  | SHAM MALE | 12.8 | 28.2 | 2.1 | 11 |  |  |
|  |  | CBD MALE | 20.2 | 32.5 | 5.1 | 10 |  |  |
|  |  | THC MALE | 20.5 | 41.6 | 2.4 | 16 |  |  |
|  | Closed<br>arms | SHAM FEMALE | 44.6 | 72.2 | 27.3 | 13 | **Interaction:** F (2, 72) = 0,6394; P=0,5306<br>**Sex:** F (1, 72) = 0,2572; P=0,6136<br>**Treatment** F (2, 72) = 0,9359; P=0,3970 | x |
|  |  | CBD FEMALE | 46.5 | 64.8 | 28.4 | 12 |  |  |
|  |  | THC FEMALE | 42.3 | 57.5 | 25.1 | 16 |  |  |
|  |  | SHAM MALE | 38.2 | 71.5 | 25.9 | 11 |  |  |
|  |  | CBD MALE | 47.8 | 79.1 | 36.9 | 10 |  |  |
|  |  | THC MALE | 44.3 | 72.9 | 31.1 | 16 |  |  |
| SAP(%) | Center | SHAM FEMALE | 2.8 | 5.2 | 0.9 | 13 | **Interaction:** F (2, 72) = 1,026; P=0,3635<br>**Sex:** F (1, 72) = 84,53; P<0,0001<br>**Treatment** F (2, 72) = 5,502; F (2, 72) = 5,502 | **FEMALE**<br>"Sham vs. CBD" P=0,3855<br>"Sham vs. THC" P=0,1254<br>"CBD vs. THC" P=0,0040<br>**MALE**<br>"Sham vs. CBD" P=0,9995<br>"Sham vs. THC" P=0,4290<br>"CBD vs. THC" P=0,4662<br>**FEMALE vs. MALE**<br>"Sham" P<0,0001<br>"CBD" P<0,0001<br>"THC" P<0,0001 |
|  |  | CBD FEMALE | 3.4 | 5.6 | 1.3 | 12 |  |  |
|  |  | THC FEMALE | 2.2 | 3.9 | 0.6 | 16 |  |  |
|  |  | SHAM MALE | 1.1 | 1.6 | 0.3 | 11 |  |  |
|  |  | CBD MALE | 0.9 | 1.9 | 0.2 | 10 |  |  |
|  |  | THC MALE | 0.6 | 2.2 | 0 | 16 |  |  |
|  | Open<br>arms | SHAM FEMALE | 4.6 | 6.9 | 0 | 13 | **Interaction:** F (2, 72) = 0,5450; P=0,5822<br>**Sex:** F (1, 72) = 30,06; P<0,0001<br>**Treatment** F (2, 72) = 3,231; P=0,0453 | **FEMALE**<br>"Sham vs. CBD" P=0,0765<br>"Sham vs. THC" P=0,3010<br>"CBD vs. THC" P=0,6691<br>**MALE**<br>"Sham vs. CBD" P=0,3507<br>"Sham vs. THC" P=0,9999<br>"CBD vs. THC" P=0,2865<br>**FEMALE vs. MALE**<br>"Sham" P=0,0192<br>"CBD" P=0,0044<br>"THC" P<0,0001 |
|  |  | CBD FEMALE | 6.4 | 8.8 | 1.6 | 12 |  |  |
|  |  | THC FEMALE | 4.7 | 10.7 | 2.4 | 16 |  |  |
|  |  | SHAM MALE | 1.9 | 5.5 | 0.9 | 11 |  |  |
|  |  | CBD MALE | 3.2 | 6.5 | 1.2 | 10 |  |  |
|  |  | THC MALE | 1.7 | 5.9 | 0.4 | 16 |  |  |
|  | Closed<br>arms | SHAM FEMALE | 5.9 | 8.9 | 3.1 | 13 | **Interaction:** F (2, 72) = 6,353; P=0,0029<br>**Sex:** F (1, 72) = 44,95; P<0,0001<br>**Treatment** F (2, 72) = 17,66; P<0,0001 | **FEMALE**<br>"Sham vs. CBD" P=0,0220<br>"Sham vs. THC" P=0,0008<br>"CBD vs. THC" P<0,0001<br>**MALE**<br>"Sham vs. CBD" P=0,7360<br>"Sham vs. THC" P=0,0687<br>"CBD vs. THC" P=0,3558<br>**FEMALE vs. MALE** |
|  |  | CBD FEMALE | 8.3 | 10.5 | 4.1 | 12 |  |  |
|  |  | THC FEMALE | 3.9 | 7.4 | 1.2 | 16 |  |  |
|  |  | SHAM MALE | 3.9 | 5.7 | 2.1 | 11 |  |  |
|  |  | CBD MALE | 3.1 | 8.1 | 1.3 | 10 |  |  |
|  |  | THC MALE | 2.6 | 5.4 | 0.6 | 16 |  | "Sham" P=0,0032<br>"CBD" P<0,0001 |
| Contracted posture(%) | Center | SHAM FEMALE | 0.06 | 0.5 | 0 | 13 | **Interaction:** F (2, 72) = 0,9436; P=0,3940<br>**Sex:** F (1, 72) = 4,565; P=0,0360<br>**Treatment** F (2, 72) = 0,6104; P=0,5459 | **FEMALE**<br>"Sham vs. CBD" P=0,2198<br>"Sham vs. THC" P=0,9226<br>"CBD vs. THC" P=0,3530<br>**MALE**<br>"Sham vs. CBD" P=0,9470<br>"Sham vs. THC" P=0,8896<br>"CBD vs. THC" P=0,9937<br>**FEMALE vs. MALE**<br>"Sham" P=0,0373<br>"CBD" P=0,8903<br>"THC" P=0,1271 |
|  |  | CBD FEMALE | 0.2 | 1.4 | 0 | 12 |  |  |
|  |  | THC FEMALE | 0.2 | 0.7 | 0 | 16 |  |  |
|  |  | SHAM MALE | 0.3 | 1.1 | 0 | 11 |  |  |
|  |  | CBD MALE | 0.2 | 1.8 | 0.02 | 10 |  |  |
|  |  | THC MALE | 0.2 | 0.9 | 0 | 16 |  |  |
|  | Open arms | SHAM FEMALE | 0 | 0.08 | 0 | 13 | **Interaction:** F (2, 68) = 0,1873; P=0,8297<br>**Sex:** F (1, 68) = 27,85; P<0,0001<br>**Treatment** F (2, 68) = 4,092; P=0,0210 | **FEMALE**<br>"Sham vs. CBD" P=0,5994<br>"Sham vs. THC" P=0,2458<br>"CBD vs. THC" P=0,8614<br>**MALE**<br>"Sham vs. CBD" P=0,2025<br>"Sham vs. THC" P=0,0530<br>"CBD vs. THC" P=0,9023<br>**FEMALE vs. MALE**<br>"Sham" P=0,0170<br>"CBD" P=0,0030<br>"THC" P=0,0003 |
|  |  | CBD FEMALE | 0.026 | 0.3 | 0 | 12 |  |  |
|  |  | THC FEMALE | 0.08 | 0.3 | 0 | 16 |  |  |
|  |  | SHAM MALE | 0.1 | 0.4 | 0 | 11 |  |  |
|  |  | CBD MALE | 0.2 | 0.5 | 0.08 | 10 |  |  |
|  |  | THC MALE | 0.3 | 0.7 | 0 | 16 |  |  |
|  | Closed arms | SHAM FEMALE | 8.4 | 22.6 | 3.7 | 13 | **Interaction:** F (2, 72) = 0,7192; P=0,4906<br>**Sex:** F (1, 72) = 0,8793; P=0,3515<br>**Treatment** F (2, 72) = 19,79; P<0,0001 | **FEMALE**<br>"Sham vs. CBD" P=0,0008<br>"Sham vs. THC" P=0,2116<br>"CBD vs. THC" P=0,0581<br>**MALE**<br>"Sham vs. CBD" P<0,0001<br>"Sham vs. THC" P=0,1856<br>"CBD vs. THC" P=0,0015<br>**FEMALE vs. MALE**<br>"Sham" P=0,2779<br>"CBD" P=0,6598<br>"THC" P=0,2756 |
|  |  | CBD FEMALE | 2.7 | 9.5 | 1.4 | 12 |  |  |
|  |  | THC FEMALE | 5.3 | 14.2 | 2.6 | 16 |  |  |
|  |  | SHAM MALE | 11.2 | 20.3 | 4.8 | 11 |  |  |
|  |  | CBD MALE | 2.1 | 5.3 | 1.1 | 10 |  |  |
|  |  | THC MALE | 8.5 | 20.2 | 3.7 | 16 |  |  |

### Prenatal cannabinoids induce a universal increase in repetitive defensive behavior

The MB task was employed as an ethological measure of defensive persistence, where the burying of non-reactive objects serves as a proxy for repetitive, stereotyped behaviors responding to potential environmental threats (22,33). This assay provides a distinct dimension of behavioral reactivity that complements the EPM by probing behavioral persistence and switching deficits. We previously reported that prenatal CBD exposure increases marble burying in both male and female offspring (15); however, the long-term impact of prenatal THC on this specific phenotype remained uncharacterized. To address this, we applied our established exposure protocol to the current cohort. Analysis revealed that offspring exposed to either CBD or THC buried significantly more marbles than Sham controls (Figure 2, Table 5). This increase was observed in both males and females, indicating a universal tendency toward behavioral perseveration associated with prenatal cannabinoid exposure. In contrast, Sham progeny exhibited comparable baseline burying levels across sexes, indicating no innate sex differences in this measure.

**Figure 2.**
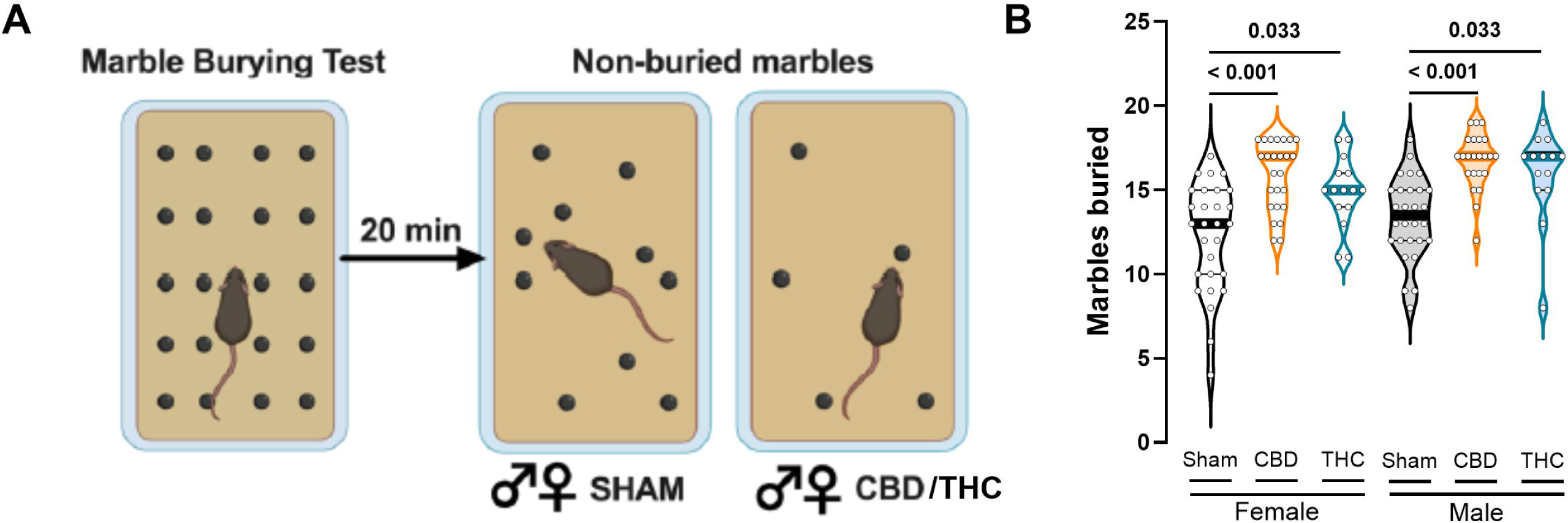
Gestational Cannabinoid Exposure Elevates Repetitive Responses in the Marble Burying Task. (A) Left: schematic representation of the marble burying paradigm. Right: representative results from Sham, CBD, and THC-exposed animals of both sexes. (B) Quantification of buried marbles revealed that both male and female offspring prenatally exposed to CBD or THC buried significantly more marbles than their Sham counterparts. Data are presented as violin plots (median, and 25th–75th percentiles), with each point representing an individual animal. Statistical analysis was performed using two-way ANOVA followed by Šídák’s multiple comparisons test. *P values < 0.05 are indicated in the graph. Group sizes were as follows: Sham female (N = 28), CBD female (N = 25), THC female (N = 15), Sham male (N = 28), CBD male (N = 23), THC male (N = 14).

**Table 5:** Analysis of MB. Statistical significance defined as p-value < 0.05.

| Measurement | Treatment | Median | Max | Min | N | Two-way ANOVA Main effects and interactions | Multiple comparison (Šidák's post hoc test) |
| --- | --- | --- | --- | --- | --- | --- | --- |
| Buried marbles | SHAM FEMALE | 13 | 17 | 4 | 28 | **Interaction:** F (2, 127) = 0,03546; P=0,9652<br>**Sex:** F (1, 127) = 5,234; P=0,0238<br>**Treatment** F (2, 127) = 28,35; P<0,0001 | **FEMALE**<br>"Sham vs. CBD" P< 0,0001<br>"Sham vs. THC" P=0,0033<br>"CBD vs. THC" P=0,5010<br>**MALE**<br>"Sham vs. CBD" P<0,0001<br>"Sham vs. THC" P=0,0050<br>"CBD vs. THC" P=0,6992<br>**FEMALE vs. MALE**<br>"Sham" P=0,0927<br>"CBD" P=0,2262<br>"THC" P=0,2448 |
|  | CBD FEMALE | 17 | 18 | 12 | 25 |  |  |
|  | THC FEMALE | 15 | 18 | 11 | 15 |  |  |
|  | SHAM MALE | 13.5 | 18 | 8 | 28 |  |  |
|  | CBD MALE | 17 | 19 | 12 | 23 |  |  |
|  | THC MALE | 17 | 19 | 8 | 14 |  |  |

### From behavioral rigidity to circuit remodeling

The convergence of motor stereotypies and prolonged risk assessment indicates a fundamental breakdown in the strategies required to navigate environmental uncertainty and switch actions (22,33). Given that the medial prefrontal cortex (mPFC) exerts top-down control over defensive behaviors (16–18), we hypothesized that these phenotypes stem from a remodeling of prefrontal efferent circuitry. To isolate the underlying cellular substrates, we interrogated Layer 5 (L5) pyramidal neurons, the primary projection node of the mPFC (34–36). Specifically, we tested whether behavioral inflexibility correlates with a collapse of synaptic adaptability and intrinsic excitability, determining if a state of persistent synaptic rigidity mirrors the loss of context-dependent behavioral modulation.

### Prenatal cannabinoids differentially reshape the intrinsic excitability of mPFC projection neurons

To determine how gestational cannabinoid exposure shapes the biophysical profile of mPFC output neurons, we assessed the passive membrane properties of adult L5 pyramidal cells (5) (Supplementary Fig. 2, Table 6). While resting membrane potential (RMP), membrane capacitance, and steady-state current-voltage (I-V) relationships remained unaltered across all cohorts, intrinsic excitability was selectively modified in a sex-dependent manner. Specifically, prenatal THC, but not CBD, significantly elevated the rheobase in female offspring, raising the depolarization threshold required for action potential initiation. In contrast, male rheobase values were entirely unaffected by either treatment. Collectively, these data demonstrate that gestational cannabinoid exposure drives a targeted, sex-specific remodeling of discrete excitability parameters rather than a global disruption of core somatic resistive or capacitive properties.

**Table 6:** Active and passive membrane properties of mPFC layer V pyramidal neurons. Statistical significance defined as p-value < 0.05.

| Measure | Treatment | Median | Max | Min | n/N | Two-way ANOVA<br>Main effects and<br>interactions | Multiple comparison<br>(Šidák's post hoc test) |
| --- | --- | --- | --- | --- | --- | --- | --- |
| Resting<br>Membrane<br>Potential<br>(mV) | SHAM FEMALE | -69.96 | -61.04 | -78.74 | 34/9 | **Interaction**<br>F (2, 178) = 3,031<br>P=0,0508<br>**Sex**<br>F (1, 178) = 0,1751<br>P=0,6761<br>**Treatment**<br>F (2, 178) = 2,008<br>P=0,1373 | **FEMALE**<br>"Sham vs. CBD" P=0,2166<br>"Sham vs. THC" P=0,3717<br>"CBD vs. THC" P=0,9702<br>**MALE**<br>"Sham vs. CBD" P=0,9432<br>"Sham vs. THC" P=0,0941<br>"CBD vs. THC" P=0,0309<br>**FEMALE vs. MALE**<br>"Sham" P=0,1760<br>"CBD" P=0,9301<br>"THC" P=0,039 |
|  | CBD FEMALE | -73.52 | -60.27 | -85.39 | 31/10 |  |  |
|  | THC FEMALE | -72.75 | -63.74 | -80.74 | 26/9 |  |  |
|  | SHAM MALE | -73.01 | -64.13 | -83.08 | 27/8 |  |  |
|  | CBD MALE | -73.78 | -58.33 | -84.03 | 34/14 |  |  |
|  | THC MALE | -70.41 | -57.31 | -81.01 | 30/8 |  |  |
| Rheobase<br>(pA) | SHAM FEMALE | 70 | 140 | 20 | 34/9 | **Interaction**<br>F (2, 175) = 1,407<br>P=0,2477<br>**Sex**<br>F (1, 175) = 0,3864<br>P=0,5350<br>**Treatment**<br>F (2, 175) = 9,386<br>P=0,0001 | **FEMALE**<br>"Sham vs. CBD" P=0,5066<br>"Sham vs. THC" P=0,0003<br>"CBD vs. THC" P=0,0157<br>**MALE**<br>"Sham vs. CBD" P=0,2259<br>"Sham vs. THC" P=0,0763<br>"CBD vs. THC" P=0,8133<br>**FEMALE vs. MALE**<br>"Sham" P=0,9764<br>"CBD" P=0,5369<br>"THC" P=0,1014 |
|  | CBD FEMALE | 70 | 170 | 10 | 30/10 |  |  |
|  | THC FEMALE | 110 | 210 | 40 | 26/9 |  |  |
|  | SHAM MALE | 60 | 170 | 30 | 27/8 |  |  |
|  | CBD MALE | 80 | 160 | 20 | 34/14 |  |  |
|  | THC MALE | 90 | 230 | 20 | 30/8 |  |  |
| Capacitance<br>(pF) | SHAM FEMALE | 126.38 | 243.34 | 64.22 | 35/9 | **Interaction**<br>F (2, 165) = 2,429<br>P=0,0912<br>**Sex**<br>F (1, 165) = 1,824<br>P=0,1787<br>**Treatment**<br>F (2, 165) = 1,970<br>P=0,1428 | **FEMALE**<br>"Sham vs. CBD" P=0,9555<br>"Sham vs. THC" P=0,1038<br>"CBD vs. THC" P=0,0673<br>**MALE**<br>"Sham vs. CBD" P=0,2672<br>"Sham vs. THC" P=0,7178<br>"CBD vs. THC" P=0,7218<br>**FEMALE vs. MALE**<br>"Sham" P=0,5480<br>"CBD" P=0,0110<br>"THC" P=0,5313 |
|  | CBD FEMALE | 136.41 | 259.52 | 45.07 | 30/10 |  |  |
|  | THC FEMALE | 106.72 | 176.69 | 29.94 | 22/9 |  |  |
|  | SHAM MALE | 131.62 | 210.33 | 61.83 | 24/8 |  |  |
|  | CBD MALE | 102.99 | 222.68 | 35.81 | 33/14 |  |  |
|  | THC MALE | 106.63 | 180.86 | 40.08 | 27/8 |  |  |
| Threshold<br>(mV) | SHAM FEMALE | -41.14 | -31.95 | -48.50 | 34/9 | **Interaction**<br>F (2, 176) = 0,9864<br>P=0,3750<br>**Sex**<br>F (1, 176) = 0,01012<br>P=0,9200<br>**Treatment** | **FEMALE**<br>"Sham vs. CBD" P=0,8112<br>"Sham vs. THC" P=0,2421<br>"CBD vs. THC" P=0,0815<br>**MALE**<br>"Sham vs. CBD" P=0,6621<br>"Sham vs. THC" P=0,4598 |
|  | CBD FEMALE | -39.92 | -32.53 | -53.62 | 31/10 |  |  |
|  | THC FEMALE | -42.40 | -29.88 | -50.00 | 26/9 |  |  |
|  | SHAM MALE | -39.95 | -34.73 | -48.18 | 27/8 | F (2, 176) = 2,382<br>P=0,0953 | "CBD vs. THC" P=0,9273<br>**FEMALE vs. MALE**<br>"Sham" P=0,6825<br>"CBD" P=0.2769 |
|  | CBD MALE | -41.03 | -36.96 | -47.87 | 34/14 |  |  |
|  | THC MALE | -42.44 | -35.46 | -48.98 | 30/8 |  |  |
| **Number of**<br>**APs (+400pA)** | SHAM FEMALE | 17 | 23 | 9 | 33/9 | **Interaction**<br>F (2, 178) = 3,031<br>P=0,0508<br>**Sex**<br>F (1, 178) = 0,1751<br>P=0,6761<br>**Treatment**<br>F (2, 178) = 2,008<br>P=0,1373 | **FEMALE**<br>"Sham vs. CBD" P=0,2166<br>"Sham vs. THC" P=0,3717<br>"CBD vs. THC" P=0,9702<br>**MALE**<br>"Sham vs. CBD" P=0,9432<br>"Sham vs. THC" P=0,0941<br>"CBD vs. THC" P=0,0309<br>**FEMALE vs. MALE**<br>"Sham" P=0,1760<br>"CBD" P=0.9301<br>"THC" P=0.039 |
|  | CBD FEMALE | 17 | 29 | 11 | 31/10 |  |  |
|  | THC FEMALE | 17 | 27 | 9 | 26/9 |  |  |
|  | SHAM MALE | 17 | 31 | 11 | 27/8 |  |  |
|  | CBD MALE | 18.5 | 35 | 7 | 34/14 |  |  |
|  | THC MALE | 21 | 50 | 9 | 30/8 |  |  |

**Table 7:** Voltage membrane response per current steps in mPFC pyramidal neurons in both sexes and treatment. Statistical significance defined as p-value < 0.05.

| Measure | Injected Current (pA) | Female Sham |  |  | Female CBD |  |  | corrected p-value<br>Multiple Mann-Whitney test | Female THC |  |  | corrected p-value<br>Multiple Mann-Whitney test |
| --- | --- | --- | --- | --- | --- | --- | --- | --- | --- | --- | --- | --- |
|  |  | Mea n | SEM | n/ N | Mea n | SEM | n/N |  | Mean | SEM | n/ N |  |
| Voltage changes per current step (ΔVm) | -400 | -39.13 | 1.66 | 33/9 | -39.84 | 2.18 | 31/10 | >0.99 | -33.44 | 1.95 | 26/9 | 0.24 |
|  | -350 | -35.88 | 1.55 |  | -36.41 | 1.98 |  | >0.99 | -30.53 | 1.81 |  | 0.24 |
|  | -300 | -32.27 | 1.42 |  | -32.81 | 1.81 |  | >0.99 | -27.26 | 1.64 |  | 0.24 |
|  | -250 | -28.72 | 1.33 |  | -28.99 | 1.64 |  | >0.99 | -23.92 | 1.50 |  | 0.21 |
|  | -200 | -24.84 | 1.21 |  | -25.02 | 1.49 |  | >0.99 | -20.22 | 1.34 |  | 0.17 |
|  | -150 | -20.44 | 1.09 |  | -20.67 | 1.34 |  | >0.99 | -16.31 | 1.15 |  | 0.17 |
|  | -100 | -15.07 | 0.87 |  | -15.34 | 1.10 |  | >0.99 | -11.70 | 0.88 |  | 0.12 |
|  | -50 | -8.52 | 0.54 |  | -8.71 | 0.73 |  | >0.99 | -6.38 | 0.55 |  | 0.11 |
|  | 0 | -0.02 | 0.07 |  | 0.01 | 0.07 |  | 1.0 | 0.04 | 0.06 |  | 0.55 |
|  | 50 | 11.61 | 1.03 |  | 10.65 | 1.00 |  | 1.0 | 8.23 | 0.90 |  | 0.17 |
|  | 100 | 19.61 | 1.22 |  | 20.66 | 1.77 |  | >0.99 | 15.81 | 1.41 |  | 0.24 |
|  | 150 | 24.51 | 1.26 |  | 26.20 | 1.68 |  | 1.0 | 22.49 | 1.91 |  | 0.55 |
|  | 200 | 31.96 | 1.85 |  | 31.50 | 1.92 |  | >0.99 | 29.07 | 2.31 |  | 0.55 |
|  | Injected Current (pA) | Male Sham |  |  | Male CBD |  |  | corrected p-value<br>Multiple Mann-Whitney test | Male THC |  |  | corrected p-value<br>Multiple Mann-Whitney test |
|  |  | Mea n | SEM | n/ N | Mea n | SEM | n/N |  | Mean | SEM | n/ N |  |
| Voltage changes per current step (ΔVm) | -400 | -46.43 | 20.91 | 27/8 | -38.23 | 11.54 | 34/14 | 0.64 | -45.41 | 16.66 | 30/8 | 1.00 |
|  | -350 | -42.01 | 17.90 |  | -34.58 | 9.75 |  | 0.64 | -41.05 | 14.28 |  | 1.00 |
|  | -300 | -37.69 | 15.08 |  | -31.05 | 8.72 |  | 0.64 | -36.65 | 12.47 |  | 1.00 |
|  | -250 | -32.86 | 12.90 |  | -27.31 | 7.73 |  | 0.64 | -31.89 | 10.63 |  | 1.00 |
|  | -200 | -28.27 | 11.09 |  | -23.48 | 6.94 |  | 0.64 | -26.99 | 8.93 |  | 1.00 |
|  | -150 | -23.42 | 9.16 |  | -19.25 | 6.00 |  | 0.64 | -21.47 | 7.23 |  | 0.99 |
|  | -100 | -17.47 | 7.28 |  | -14.25 | 4.75 |  | 0.64 | -15.48 | 5.51 |  | 0.98 |
|  | -50 | -10.04 | 4.39 |  | -7.89 | 3.10 |  | 0.64 | -8.52 | 3.17 |  | 0.91 |
|  | 0 | 0.15 | 0.43 |  | 0.07 | 0.53 |  | 0.77 | -0.05 | 0.44 |  | 0.83 |
|  | 50 | 12.86 | 6.53 |  | 10.49 | 4.26 |  | 0.74 | 9.53 | 3.33 |  | 0.45 |
|  | 100 | 22.12 | 9.56 |  | 20.46 | 8.60 |  | 0.77 | 16.69 | 5.92 |  | 0.08 |
|  | 150 | 26.87 | 7.97 |  | 26.15 | 6.81 |  | 0.77 | 23.01 | 8.55 |  | 0.31 |
|  | 200 | 32.32 | 8.79 |  | 30.97 | 8.60 |  | 0.75 | 24.82 | 6.98 |  | 0.01 |

### Prenatal THC augments the repetitive firing capacity of male mPFC projection neurons

Active firing dynamics were assessed via current–spike frequency (I-F) curves in adult L5 pyramidal neurons (Figure 3, Table 8). Paralleling prior rodent data (5), prenatal THC selectively disrupted intrinsic excitability in male offspring. Female cohorts displayed conserved firing rates across all treatment conditions (Figure 3A). In contrast, THC-exposed males exhibited a robust potentiation of firing capacity, characterized by significantly elevated spike frequencies during depolarizing current injections, particularly at higher stimulus intensities (Figure 3B–D). This hyperexcitability developed independently of changes to the spike-initiation gate, as action potential (AP) thresholds remained constant across all sexes and treatments (Figure 3C). Together, these results reveal that prenatal THC exposure persistently recalibrates male mPFC intrinsic excitability without altering baseline spike generation mechanics.

**Figure 3.**
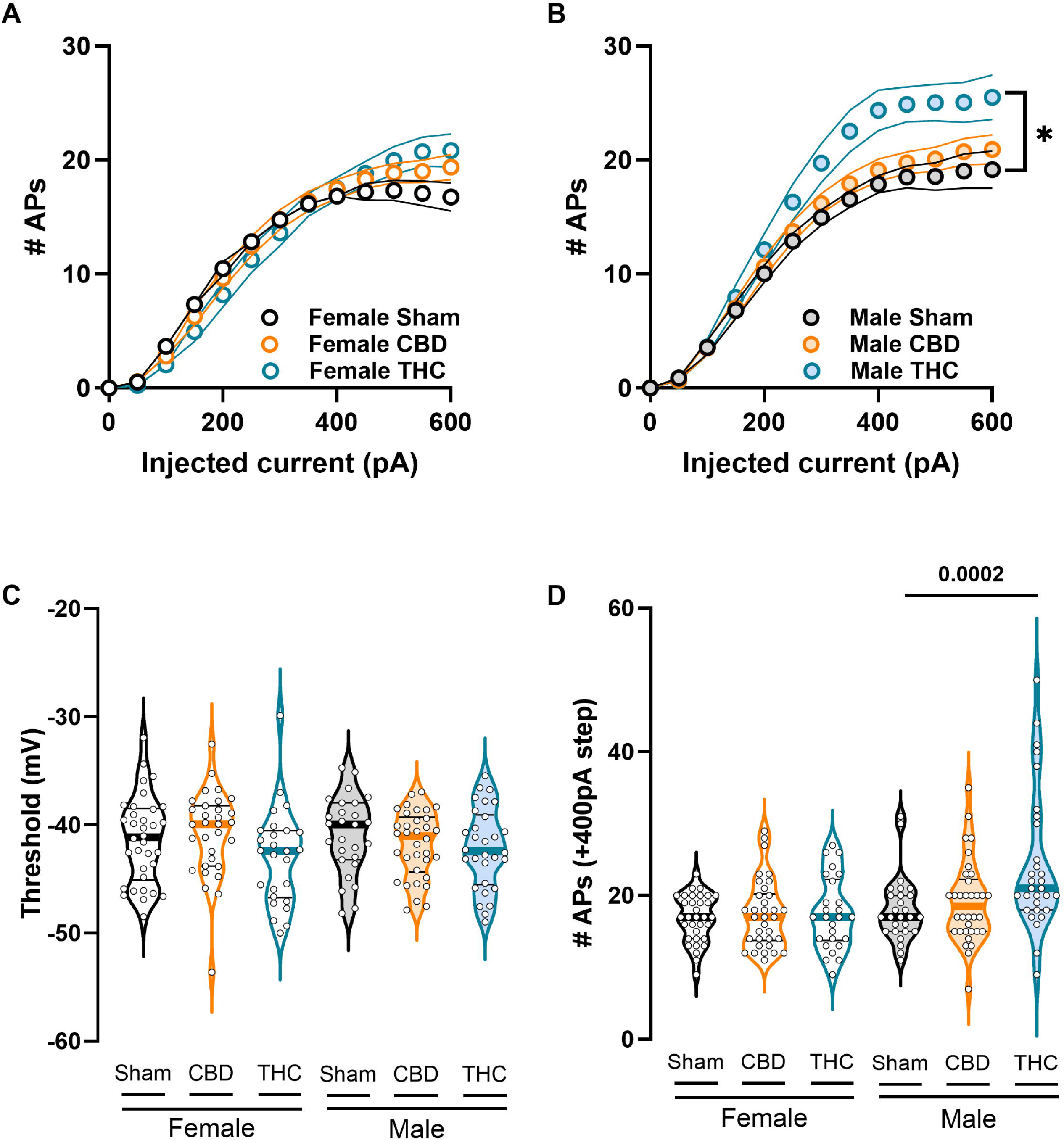
THC-exposed male progeny displays increased intrinsic neuronal excitability in adulthood. (A–B) Number of evoked action potentials in response to increasing depolarizing current injections. (A) Female offspring display comparable intrinsic excitability across experimental conditions. (B) Progressive depolarizing current injections (50-pA steps) reveal increased firing in THC-exposed males at supraphysiological stimulation intensities. (C) Despite this increase in firing, the action potential threshold does not differ across sex or treatment. (D) Representative example showing the number of spikes elicited by a +400-pA depolarizing current step, illustrating enhanced firing in THC-exposed males. (A–B) Each dot represents the group mean value at the corresponding current step; data are presented as mean ± SEM in XY plots and were analyzed using multiple measures Mann-Whitney Test. (C–D) Each dot represents a single neuron; data are shown as violin plots (median, and 25th–75th percentiles) and were analyzed using two-way ANOVA followed by Šídák’s multiple-comparison test. Only statistically significant differences (p < 0.05) are indicated in the graphs. Sample sizes (# neurons / # animals) were females Sham (34/9), CBD (31/10), THC (26/9); males Sham (27/8), CBD (34/14), THC (30/8).

**Table 8:**
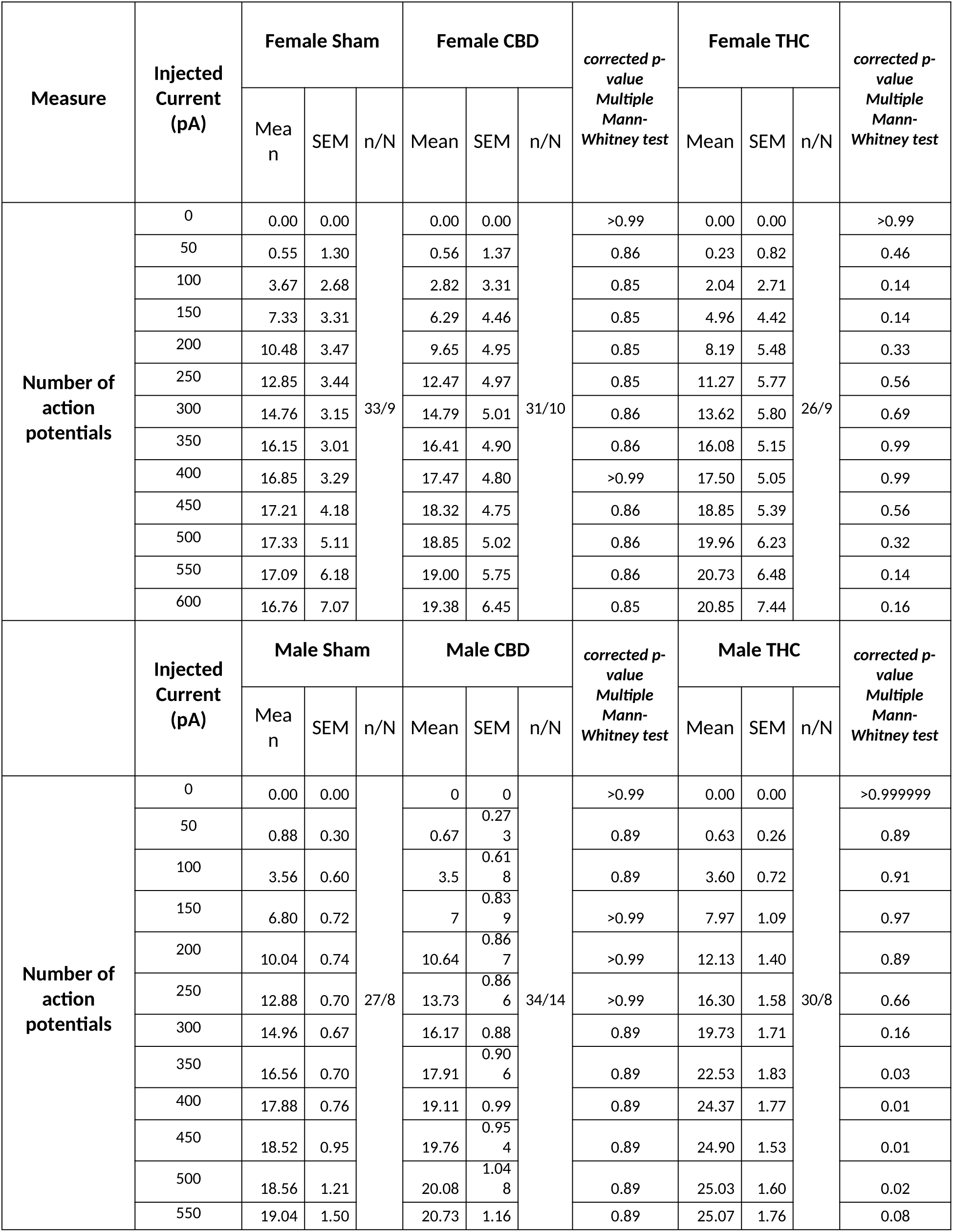
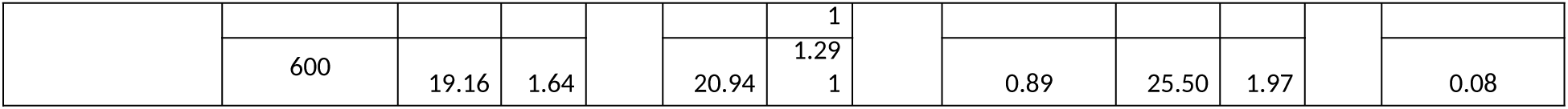
Voltage membrane response per current steps in mPFC pyramidal neurons in both sexes and treatment. Statistical significance defined as p-value < 0.05.

### Prenatal cannabinoids are associated with sex-divergent rescaling of excitatory drive and synaptic kinetics

To determine if altered intrinsic excitability correlates with reshaped synaptic inputs, we recorded spontaneous excitatory postsynaptic currents (sEPSCs) in L5 pyramidal neurons (Figure 4, Table 9). In female offspring, prenatal CBD robustly enhanced excitatory drive, significantly increasing mean sEPSC amplitude and frequency (Supplementary Fig. 3A-B). Log-normal analysis revealed a categorical rightward shift toward high-amplitude events and a concurrent shortening of inter-event intervals, indicating an elevated glutamate release rate (Figure 4C,E). Prenatal THC similarly shifted the female amplitude distribution rightward without altering global means, suggesting a more restricted remodeling of the synaptic pool. Male offspring displayed a distinct phenotype; CBD-exposed males exhibited subtle distributional shifts toward small-amplitude events and shortened inter-event intervals that failed to alter global means. Beyond event frequency and magnitude, prenatal THC accelerated sEPSC rise-time kinetics across both sexes while decay times remained stable (Figure 4H-I). This selective abbreviation of the rising phase points to discrete remodeling of the synaptic microarchitecture, potentially reflecting altered AMPA receptor subunit stoichiometry or a proximal redistribution of excitatory inputs.

**Figure 4.**
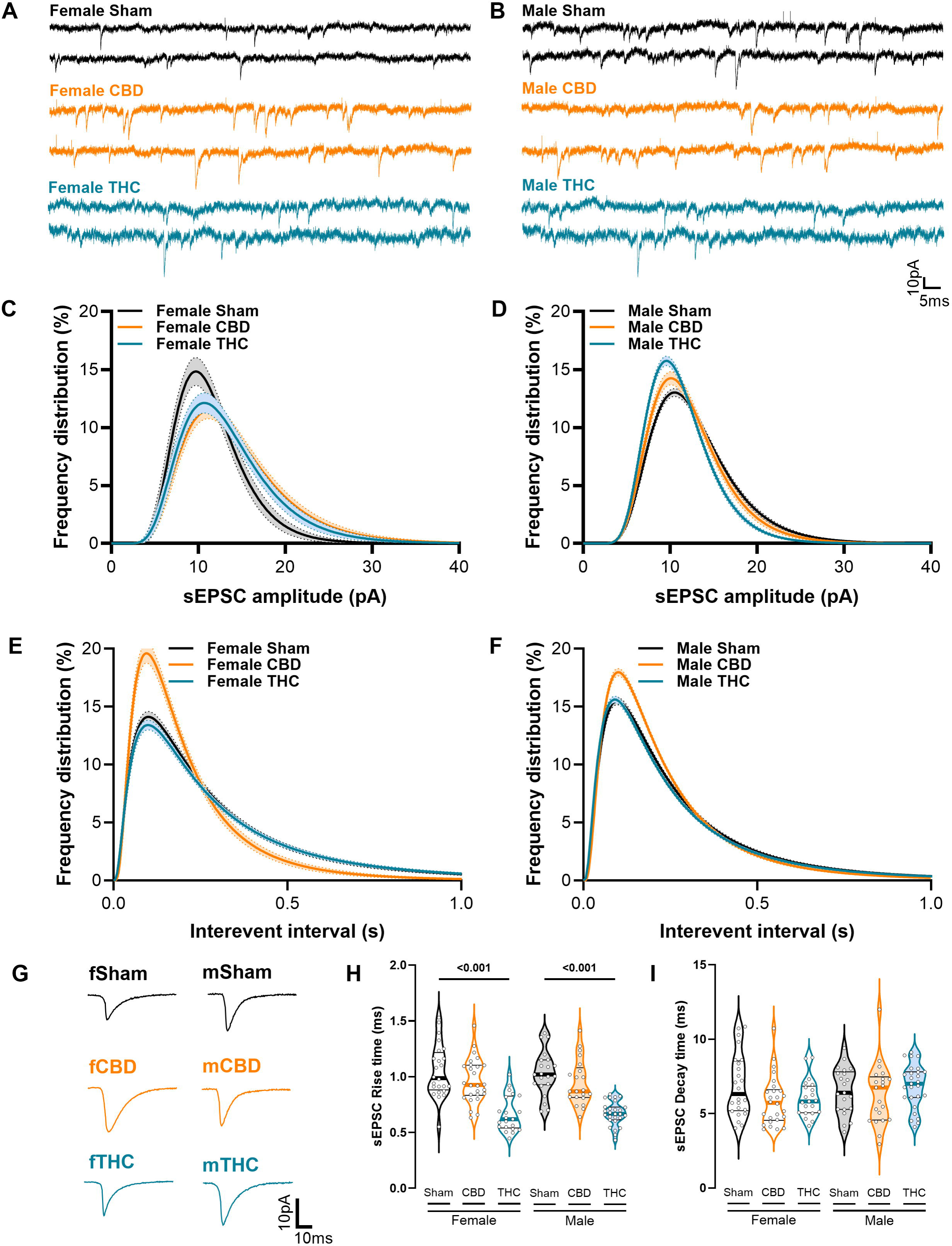
Prenatal cannabinoid exposure alters excitatory synaptic transmission in female progeny: (A–B) Representative traces of spontaneous excitatory postsynaptic currents (sEPSCs) recorded at −70 mV in presence of gabazine (10µM) from Sham-, CBD-, and THC-exposed female and male offspring (scale bar 10pA/5ms). (C) Log-normal distribution fitting (± CI) reveals a higher proportion of large-amplitude events in cannabinoid-treated females, whereas (D) cannabinoid-treated males display a shift toward smaller-amplitude events. (E) IInter-event intervals are reduced in CBD-exposed females relative to Sham and THC females. (F) A higher proportion of CBD-exposed males exhibit shorter inter-event intervals, indicating increased event frequency. (G) Representative traces of averaged excitatory transmission currents. (H) Regardless of sex, THC-exposed progeny exhibits faster rise times of excitatory postsynaptic currents. (I) Decay times of excitatory postsynaptic currents remain unchanged across sex and treatment. (C-F) Data were analyzed using log-normal curve fitting with confidence intervals (± CI). (H, I) Data are presented as violin plots (median, and 25th–75th percentiles) and were analyzed using two-way ANOVA followed by Šídák’s correction for multiple comparisons. Only statistically significant differences (p < 0.05) are indicated in the graphs additional statistical information can be found in Table 9. Sample sizes (# neurons / # animals) were females Sham (24/9), CBD (26/12), THC (21/9); males Sham (21/8), CBD (21/13), THC (27/8).

**Table 9:** Characterization of excitatory postsynaptic currents features recorded in layer V pyramidal neurons. Statistical significance defined as p-value < 0.05.

| Measure | Sex | Treatment | Median | MAX | MIN | n/N | Two-way ANOVA Main effects and interactions | Multiple comparison (Šidák's post hoc test) |
| --- | --- | --- | --- | --- | --- | --- | --- | --- |
| sEPSCs Amplitude (pA) | Female | Sham | 13.94 | 23.13 | 10.31 | 24//9 | **Interaction**<br>F (2, 178) = 3,031<br>P=0,0508<br>**Sex**<br>F (1, 178) = 0,1751 P=0,6761<br>**Treatment**<br>F (2, 178) = 2,008<br>P=0,1373 | **FEMALE**<br>"Sham vs. CBD" P=0,0338<br>"Sham vs. THC" P=0,0596<br>"CBD vs. THC" P=0,9945<br>**MALE**<br>"Sham vs. CBD" P=0,5135<br>"Sham vs. THC" P=0,1286<br>"CBD vs. THC" P=0,7157<br>**FEMALE vs. MALE**<br>"Sham" P=0,0962<br>"CBD" P=0,0594<br>"THC" P=0,0105 |
|  |  | CBD | 16.47 | 23.99 | 9.03 | 26//12 |  |  |
|  |  | THC | 16.01 | 25.00 | 10.58 | 21//9 |  |  |
|  | Male | Sham | 16.13 | 23.57 | 10.60 | 21//8 |  |  |
|  |  | CBD | 13.63 | 23.86 | 10.52 | 21//13 |  |  |
|  |  | THC | 13.10 | 20.30 | 10.55 | 27//8 |  |  |
| sEPSCs Frequency (Hz) | Female | Sham | 2.64 | 6.52 | 1.23 | 24//9 | **Interaction**<br>F (2, 133) = 0,1497<br>P=0,8611<br>**Sex**<br>F (1, 133) = 0,6387<br>P=0,4256<br>**Treatment**<br>F (2, 133) = 11,17<br>P<0,0001 | **FEMALE**<br>"Sham vs. CBD" P=0,0226<br>"Sham vs. THC" P=0,7347<br>"CBD vs. THC" P=0,0030<br>**MALE**<br>"Sham vs. CBD" P=0,2004<br>"Sham vs. THC" P=0,3718<br>"CBD vs. THC" P=0,0046<br>**FEMALE vs. MALE**<br>"Sham" P=0,9857<br>"CBD" P=0,4446<br>"THC" P=0,5394 |
|  |  | CBD | 4.16 | 7.49 | 1.17 | 26//12 |  |  |
|  |  | THC | 2.79 | 6.16 | 0.40 | 21//9 |  |  |
|  | Male | Sham | 2.79 | 5.38 | 1.11 | 21//8 |  |  |
|  |  | CBD | 3.74 | 6.08 | 2.07 | 21//13 |  |  |
|  |  | THC | 2.24 | 6.17 | 0.52 | 27//8 |  |  |
| Rise time (ms) | Female | Sham | 0.99 | 1.53 | 0.55 | 24//9 | **Interaction**<br>F (2, 134) = 0,06577<br>P=0,9364<br>**Sex**<br>F (1, 134) = 0,1151<br>P=0,7349<br>**Treatment**<br>F (2, 134) = 50,47<br>P<0,0001 | **FEMALE**<br>"Sham vs. CBD" P=0,2539<br>"Sham vs. THC" P=<0,0001<br>"CBD vs. THC" P=<0,0001<br>**MALE**<br>"Sham vs. CBD" P=0,2956<br>"Sham vs. THC" P=<0,0001<br>"CBD vs. THC" P=<0,0001<br>**FEMALE vs. MALE**<br>"Sham" P=0,7517<br>"CBD" P=0,7161<br>"THC" P=0,9217 |
|  |  | CBD | 0.93 | 1.46 | 0.63 | 26//12 |  |  |
|  |  | THC | 0.62 | 1.02 | 0.45 | 21//9 |  |  |
|  | Male | Sham | 1.02 | 1.39 | 0.69 | 21//8 |  |  |
|  |  | CBD | 0.87 | 1.41 | 0.64 | 21//13 |  |  |
|  |  | THC | 0.67 | 0.84 | 0.43 | 27//8 |  |  |
| Decay time (ms) | Female | Sham | 6.32 | 10.85 | 4.03 | 24//9 | **Interaction**<br>F (2, 134) = 1,447<br>P=0,2390<br>**Sex**<br>F (1, 134) = 1,087<br>P=0,2990<br>**Treatment** | **FEMALE**<br>"Sham vs. CBD" P=0,0887<br>"Sham vs. THC" P=0,2037<br>"CBD vs. THC" P=0,9503<br>**MALE**<br>"Sham vs. CBD" P=0,9136<br>"Sham vs. THC" P=0,8032 |
|  |  | CBD | 5.73 | 10.73 | 3.95 | 26//12 |  |  |
|  |  | THC | 5.82 | 8.77 | 4.18 | 21//9 |  |  |
|  | Male | Sham | 6.40 | 9.41 | 3.76 | 21//8 | F (2, 134) = 1,511<br>P=0,2244 | "CBD vs. THC" P=0,5401<br>**FEMALE vs. MALE**<br>"Sham" P=0,4761<br>"CBD" P=0,3718 |
|  |  | CBD | 6.75 | 12.00 | 2.95 | 21//13 |  |  |
|  |  | THC | 7.01 | 9.12 | 4.11 | 27//8 |  |  |

### Prenatal cannabinoids are associated with sex-specific remodeling of the inhibitory landscape and temporal tuning

To determine if the GABAergic microcircuitry underwent a concomitant reorganization, we recorded spontaneous inhibitory postsynaptic currents (sIPSCs) in adult L5 pyramidal neurons (Figure 5, Table 10). In female offspring, prenatal CBD robustly enhanced inhibitory transmission, significantly elevating mean sIPSC amplitudes (Supplementary Fig. 4A) and enriching large-amplitude events (Figure 5C) at a stable frequency. Conversely, THC-exposed females exhibited an attenuated profile; while average frequency was conserved, distributional analysis revealed an increased proportion of longer inter-event intervals, indicating reduced GABA_A_ receptor activation probability (Figure 5E-F). In male offspring, gestational cannabinoids generally weakened inhibitory strength: prenatal THC significantly decreased mean amplitudes (Supplementary Fig. 4A), and both compounds shifted event distributions toward smaller amplitudes (Figure 5D) without modifying average frequencies.

**Figure 5.**
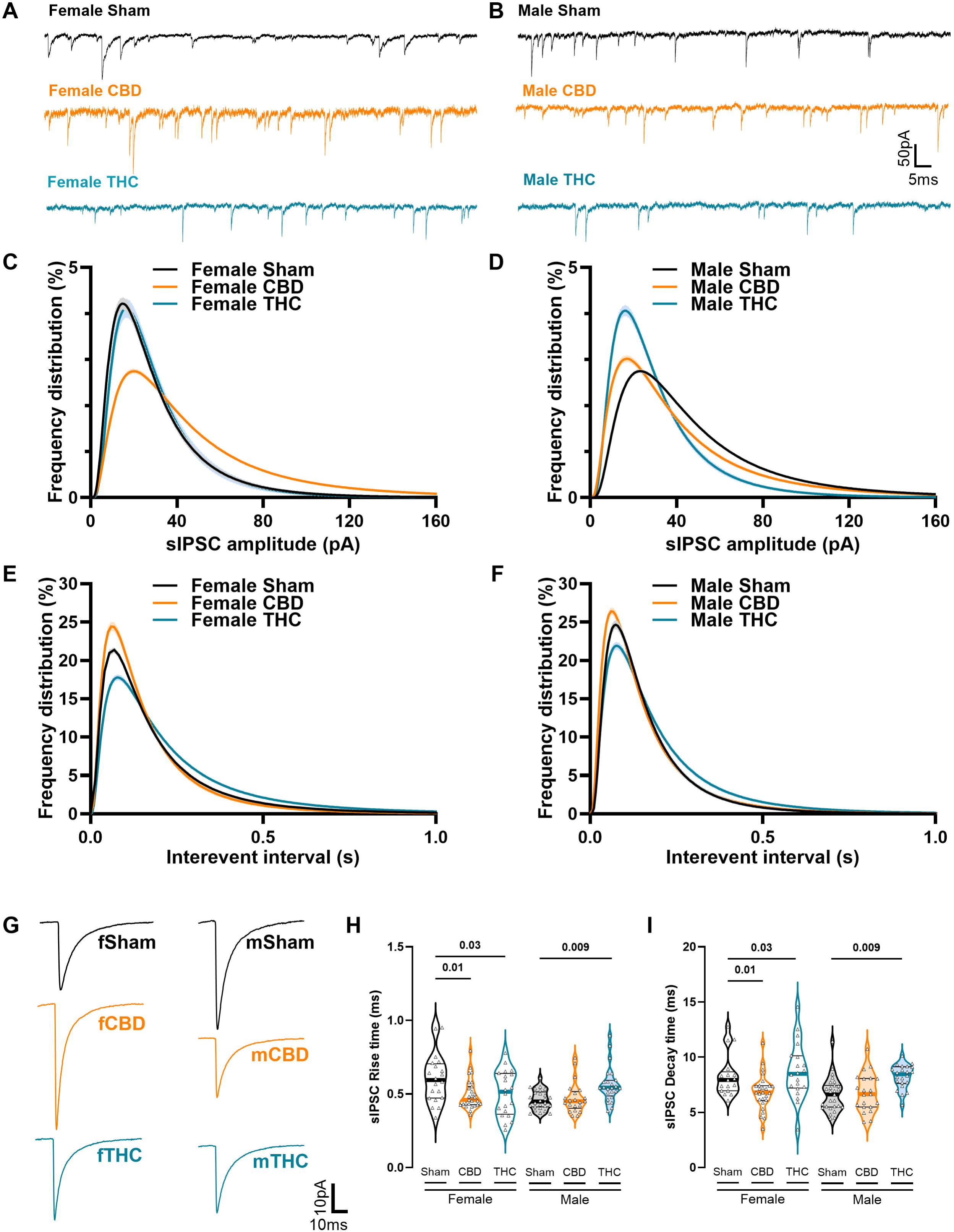
Prenatal cannabinoid exposure differentially alters inhibitory synaptic transmission in female and male progeny: (A–B) Representative traces of spontaneous inhibitory postsynaptic currents (sIPSCs) recorded at −70 mV from Sham-, CBD-, and THC-exposed female and male offspring in the presence of APV and CNQX (10 µM), (scale bars: 50 pA, 5 ms). (C) Log-normal distribution fitting (± CI) reveals a higher proportion of large-amplitude events in CBD-treated females, whereas (D) cannabinoid-treated males display a shift toward smaller-amplitude events. (E) In the distributions, inter-event intervals are reduced in CBD-exposed females relative to Sham and THC females. (F) Conversely, THC-exposed males display a higher proportion of longer inter-event intervals, indicative of reduced event frequency. (G) Representation on mean sIPSC currents per sex and treatment. (H) Compared with controls, inhibitory postsynaptic currents display faster rise times in cannabinoid-treated females, whereas cannabinoid-treated males exhibit slower rise times. (I) For inhibitory current decay kinetics, CBD-exposed females show faster decay times, while THC-exposed females exhibit prolonged decays; THC-exposed males also display increased decay times. (C-F) The graphs display the relative distribution of all events during 6 min recording, these data were analyzed using log-normal curve fitting with confidence intervals (± CI). (H, I) Data are presented as violin plots (median, and 25th–75th percentiles) and were analyzed using two-way ANOVA followed by Šídák’s correction for multiple comparisons. Only statistically significant differences (p < 0.05) are indicated in the graphs. Further statistical details can be found on Table 6. Sample sizes (# neurons / # animals) were females Sham (20/6), CBD (27/7), THC (21/5); males Sham (27/7), CBD (21/7), THC (24/5).

**Table 10:** Characterization of inhibitory postsynaptic currents features recorded in layer V pyramidal neurons. Statistical significance defined as p-value < 0.05.

| Measure | Sex | Treatment | Median | MAX | MIN | n/N | Two-way ANOVA<br>Main effects and<br>interactions | Multiple comparison<br>(Šidák's post hoc test) |
| --- | --- | --- | --- | --- | --- | --- | --- | --- |
| sIPSCs<br>Amplitude<br>(pA) | Female | Sham | 33.62 | 70.74 | 21.87 | 20//6 | **Interaction**<br>F (2, 133) = 2,858<br>P=0,0609<br>**Sex**<br>F (1, 133) = 1,456<br>P=0,2297<br>**Treatment**<br>F (2, 133) = 8,210<br>P=0,0004 | **FEMALE**<br>"Sham vs. CBD" P=0,0219<br>"Sham vs. THC" P=0,9911<br>"CBD vs. THC" P=0,0150<br>**MALE**<br>"Sham vs. CBD" P=0,9796<br>"Sham vs. THC" P=0,0037<br>"CBD vs. THC" P=0,0122<br>**FEMALE vs. MALE**<br>"Sham" P=0,0089<br>"CBD" P=0,8213<br>"THC" P=0,7457 |
|  |  | CBD | 55.64 | 86.65 | 14.58 | 27//7 |  |  |
|  |  | THC | 36.46 | 76.54 | 18.34 | 21//5 |  |  |
|  | Male | Sham | 51.75 | 75.86 | 29.07 | 27//7 |  |  |
|  |  | CBD | 48.26 | 105.22 | 10.33 | 21//7 |  |  |
|  |  | THC | 36.51 | 56.71 | 15.67 | 24//5 |  |  |
| sIPSCs<br>Frequency<br>(Hz) | Female | Sham | 5.31 | 8.66 | 0.35 | 20//6 | **Interaction**<br>F (2, 133) = 0,3607<br>P=0,6979<br>**Sex**<br>F (1, 133) = 3,416<br>P=0,0668<br>**Treatment**<br>F (2, 133) = 4,351<br>P=0,0148 | **FEMALE**<br>"Sham vs. CBD" P=0,3179<br>"Sham vs. THC" P=0,4901<br>"CBD vs. THC" P=0,0227<br>**MALE**<br>"Sham vs. CBD" P=0,8261<br>"Sham vs. THC" P=0,5837<br>"CBD vs. THC" P=0,2922<br>**FEMALE vs. MALE**<br>"Sham" P=0,2071<br>"CBD" P=0,6873<br>"THC" P=0,1335 |
|  |  | CBD | 5.63 | 13.24 | 0.66 | 27//7 |  |  |
|  |  | THC | 4.01 | 8.76 | 1.02 | 21//5 |  |  |
|  | Male | Sham | 6.12 | 9.41 | 0.73 | 27//7 |  |  |
|  |  | CBD | 6.84 | 11.07 | 1.52 | 21//7 |  |  |
|  |  | THC | 4.95 | 10.34 | 1.59 | 24//5 |  |  |
| Rise time<br>(ms) | Female | Sham | 0.59 | 0.95 | 0.34 | 20//6 | **Interaction**<br>F (2, 132) = 7,901<br>P=0,0006<br>**Sex**<br>F (1, 132) = 2,956<br>P=0,0879<br>**Treatment**<br>F (2, 132) = 2,282<br>P=0,1061 | **FEMALE**<br>"Sham vs. CBD" P=0,0102<br>"Sham vs. THC" P=0,0281<br>"CBD vs. THC" P=0,9834<br>**MALE**<br>"Sham vs. CBD" P=0,9120<br>"Sham vs. THC" P=0,0086<br>"CBD vs. THC" P=0,0404<br>**FEMALE vs. MALE**<br>"Sham" P=0,0001<br>"CBD" P=0,5114<br>"THC" P=0,1072 |
|  |  | CBD | 0.46 | 0.80 | 0.36 | 27//7 |  |  |
|  |  | THC | 0.52 | 0.78 | 0.26 | 21//5 |  |  |
|  | Male | Sham | 0.45 | 0.61 | 0.37 | 27//7 |  |  |
|  |  | CBD | 0.45 | 0.75 | 0.35 | 21//7 |  |  |
|  |  | THC | 0.54 | 0.90 | 0.38 | 24//5 |  |  |
| Decay time<br>(ms) | Female | Sham | 7.93 | 12.75 | 6.08 | 20//6 | **Interaction**<br>F (2, 127) = 2,480<br>P=0,0878<br>**Sex**<br>F (1, 127) = 5,846<br>P=0,0170<br>**Treatment**<br>F (2, 127) = 9,965<br>P<0,0001 | **FEMALE**<br>"Sham vs. CBD" P=0,0228<br>"Sham vs. THC" P=0,7587<br>"CBD vs. THC" P=0,0013<br>**MALE**<br>"Sham vs. CBD" P=0,9496<br>"Sham vs. THC" P=0,0052<br>"CBD vs. THC" P=0,0210<br>**FEMALE vs. MALE**<br>"Sham" P=0,0024 |
|  |  | CBD | 6.81 | 11.29 | 3.52 | 27//7 |  |  |
|  |  | THC | 8.49 | 14.56 | 3.45 | 21//5 |  |  |
|  | Male | Sham | 6.63 | 11.42 | 4.45 | 27//7 |  |  |
|  |  | CBD | 6.65 | 10.74 | 4.13 | 21//7 |  | "CBD" P=0,9041 |
|  |  | THC | 8.45 | 10.12 | 5.67 | 24//5 |  | "THC" P=0,3689 |

The temporal precision of inhibition was also differentially modulated by compound and sex (Figure 5H-I). In females, both cannabinoids accelerated sIPSC rise times, but their effects on decay kinetics diverged sharply: CBD accelerated, whereas THC prolonged, the decay phase. In males, CBD had no discernable kinetic effects, whereas THC slowed inhibitory signaling by increasing both rise and decay times. Collectively, these data reveal a sexually dimorphic signature where female CBD exposure potentiates inhibitory output, while prenatal THC (in both sexes) and male CBD exposure attenuate and kinetically disrupt prefrontal inhibitory drive.

### Prenatal cannabinoids are associated with sex-specific homeostatic vs. altered rescaling of the mPFC E/I landscape

To characterize the integrated functional impact of prenatal cannabinoid exposure, we quantified the net excitatory-inhibitory (E/I) balance in adult L5 pyramidal neurons by analyzing the cumulative distributions of total synaptic charge transfer (Figure 6, Table 11). In females, prenatal CBD induced a robust rightward shift in excitatory charge (Figure 6A), paralleled by a non-significant rightward trend in inhibitory charge (Figure 6B), suggesting homeostatic rescaling. Conversely, prenatal THC drove a clear pro-excitatory shift, coupling a modest increase in excitation (Figure 6A) with a significant leftward collapse of inhibitory charge (Figure 6B). Male offspring exhibited relative synaptic resilience: CBD cohorts remained stable, while THC-exposed males showed a restricted leftward shift in inhibitory charge with unaltered excitation (Figures 6E-F).

**Figure 6.**
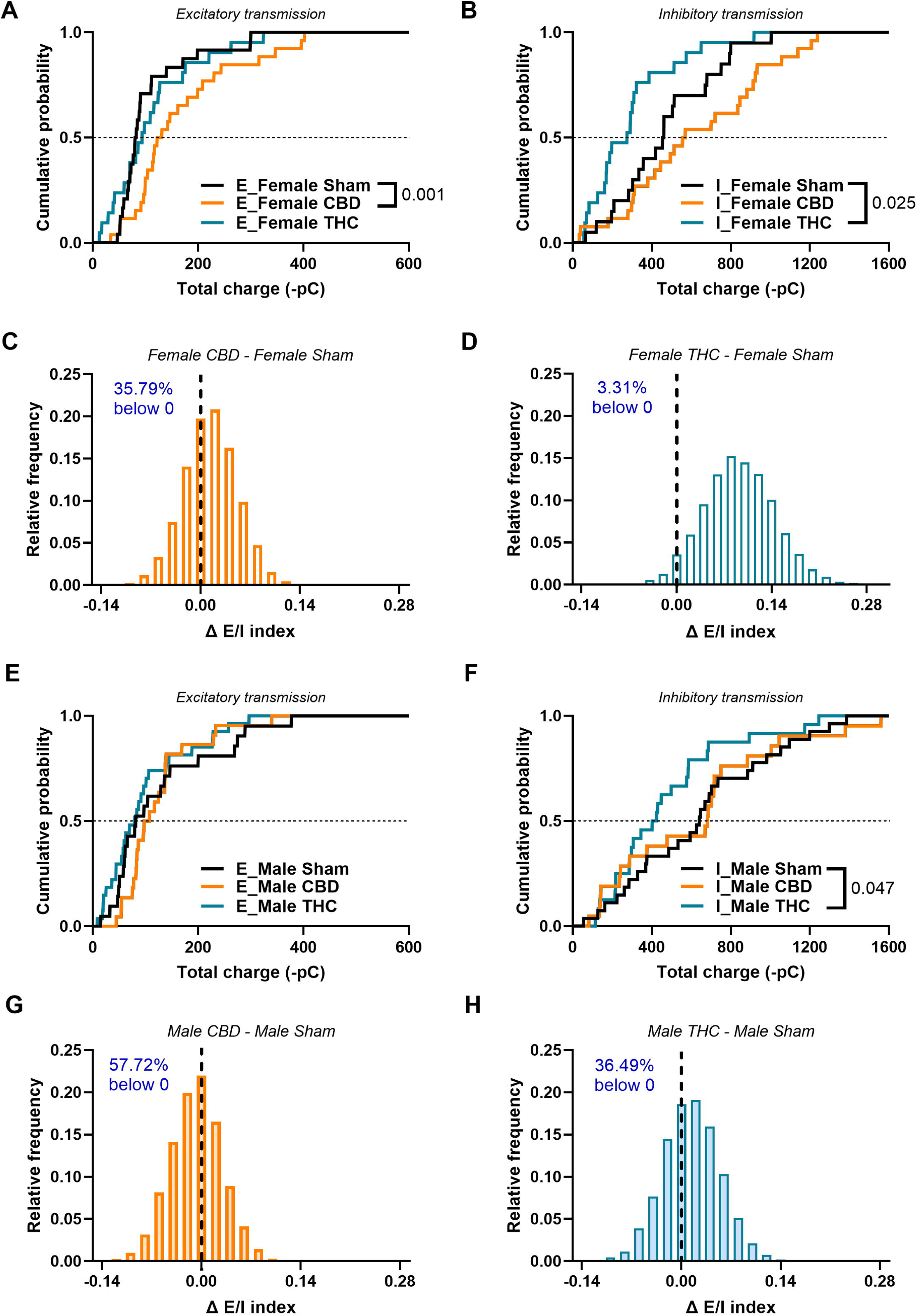
Prenatal cannabinoid exposure alters the excitatory/inhibitory balance. (A, E) Total charge transferred by AMPA-mediated sEPSCs and (B, F) GABA-mediated sIPSCs measured over a 6-min recording period across sexes and treatments. (A) CBD-exposed females show increased quantal excitatory transmission. (B) THC-exposed females display reduced inhibitory charge transfer compared with Sham controls. (C, D, G, H) Distribution plots of the difference between the bootstrapped E/I index of each cannabinoid-treated group and its corresponding Sham group. Distributions in which more than 95% of ΔE/I values fall on one side of zero are interpreted as significantly different (p < 0.05), indicating an E/I imbalance. (C) CBD-treated females show no change in E/I balance. (D) THC-treated females exhibit an increased E/I index driven by reduced inhibitory tone. (E) Total excitatory transmission remains unchanged in males, whereas (F) THC-exposed males show a marked reduction in inhibitory charge transfer. These modest synaptic changes account for the stability of the E/I index in cannabinoid-treated male offspring. (G, H) Neither CBD- nor THC-exposed males differ from Sham in their E/I index distributions. (A, B, E, F) Cumulative frequency distributions of total charge transfer for sEPSCs (excitatory) and sIPSCs (inhibitory) were obtained from the neurons recorded in Figures 4 and 5. (C, D, G, H) Relative distribution plots show the difference between the bootstrapped E/I index of each treated group and its corresponding Sham group. Only statistically significant differences (p < 0.05) are indicated in the graphs. Further statistical details on the comparisons between cumulative distributions can be found on Table 11.

**Table 11:** Comparison of cumulative distributions of excitatory/inhibitory total charge transferred by postsynaptic currents recorded in layer V pyramidal neurons. Statistical significance defined as p-value < 0.05.

| Measure | Sex | Treatment | Median | MAX | MIN | n/N | Kolmogorov-Smirnov D | Kolmogorov-Smirnov test (p-value) |
| --- | --- | --- | --- | --- | --- | --- | --- | --- |
| Total excitatory charge transferred (-pC) | Female | Sham | 81.24 | 300.00 | 46.70 | 24//9 |  |  |
|  |  | CBD | 129.40 | 402.10 | 34.22 | 26//12 | 0.555 | 0.001 |
|  |  | THC | 93.66 | 323.80 | 12.34 | 21//9 | 0.238 | 0.549 |
|  | Male | Sham | 81.49 | 376.80 | 14.82 | 21//8 |  |  |
|  |  | CBD | 102.40 | 339.50 | 44.32 | 21//13 | 0.297 | 0.301 |
|  |  | THC | 79.38 | 296.60 | 8.40 | 27//8 | 0.201 | 0.726 |
| Total inhibitory charge transferred (-pC) | Female | Sham | 458.00 | 1005.00 | 65.73 | 20//6 |  |  |
|  |  | CBD | 570.20 | 1755.00 | 33.71 | 27//7 | 0.357 | 0.106 |
|  |  | THC | 274.80 | 917.50 | 56.35 | 21//5 | 0.462 | 0.025 |
|  | Male | Sham | 640.90 | 1387.00 | 55.48 | 27//7 |  |  |
|  |  | CBD | 683.00 | 1561.00 | 81.56 | 21//7 | 0.164 | 0.908 |
|  |  | THC | 413.70 | 1246.00 | 114.40 | 24//5 | 0.384 | 0.047 |

To evaluate these shifts, we utilized a bootstrap-based estimation of the E/(E+I) index, defining significance where more than 95% of bootstrap samples fell on one side of the zero-null (Figure 6C-D, G-H). This analysis confirmed that THC-exposed females exhibit a significantly elevated E/I index, reflecting a loss of inhibitory governance (Figure 6DC). In contrast, CBD-exposed females maintained a stable net E/I balance due to parallel enhancements in both inputs (Figure 6C). Neither cannabinoid altered the net E/I index in males (Figures 6G-H). Collectively, these findings demonstrate that gestational cannabinoids reshape the mPFC along sex- and compound-specific trajectories, driving a pro-excitatory reorganization in females while sparing net synaptic balance in males.

### Prenatal CBD and THC exposure converge to abrogate eCB-LTD in the adult mPFC

To evaluate whether gestational cannabinoid exposure disrupts synaptic flexibility, we examined endocannabinoid-mediated long-term depression (eCB-LTD) in the mPFC, a plasticity hallmark vulnerable to prenatal THC in rats (5,9,10). Low-frequency stimulation (10 Hz, 10 min) induced robust, sustained eCB-LTD in Sham cohorts of both sexes (Figure 7A-D). Crucially, this form of plasticity was entirely abolished in adult mice prenatally exposed to either CBD or THC; in both sexes, fEPSP amplitudes failed to depress and remained at or above baseline (Figure 7A-D). The loss of eCB-LTD in THC-exposed female mice marks a notable species divergence from rats, where female prefrontal plasticity was previously spared (5). In this mouse model, CBD matched the efficacy of THC in nullifying eCB-LTD, indicating no protective effect. Collectively, these data demonstrate that gestational exposure to either cannabinoid converges on a shared prefrontal phenotype: the complete collapse of activity-dependent synaptic depression across both sexes (Figure 7E-F).

**Figure 7.**
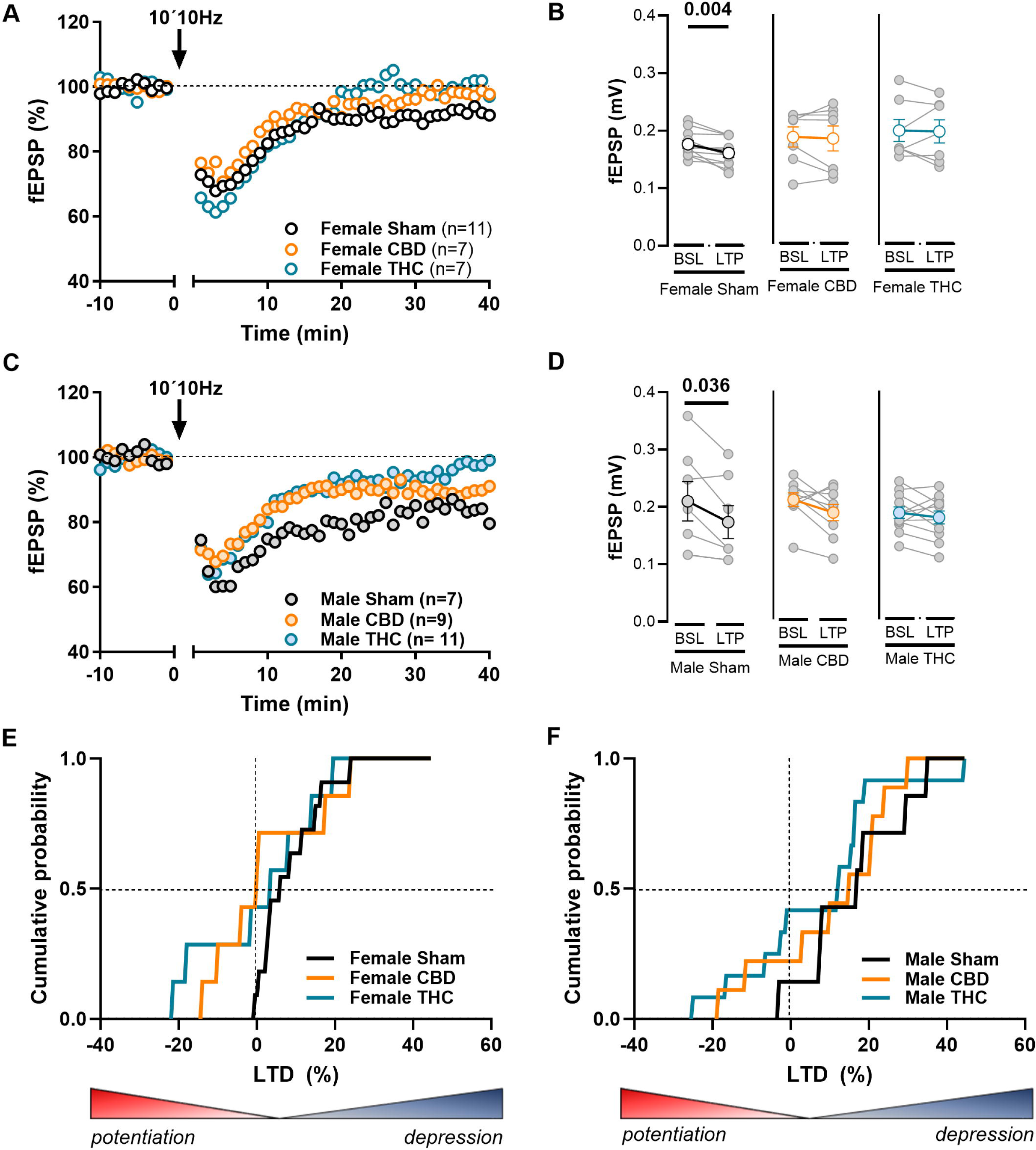
Prenatal cannabinoid exposure abolishes endocannabinoid-mediated long-term depression (eCBLTD). (A–C) Representative time courses of mean fEPSP amplitude before and after tetanic stimulation (10 × 10 Hz; arrow). (B–D) Individual experiments (grey dots) and group averages (colored circles) showing absolute fEPSP amplitudes during baseline (BSL, –10 to 0 min) and after stimulation (LTD, 30–40 min post-stimulation). Tetanic stimulation induces robust eLTD in (B) Sham females and (D) Sham males, whereas this form of synaptic plasticity is completely abolished in adult offspring prenatally exposed to CBD or THC. (E, F) Under control conditions, LTD is reliably induced in both sexes. In contrast, prenatal cannabinoid exposure abolishes LTD, with a substantial proportion of offspring instead exhibiting a potentiation response. Data are presented as mean ± SEM and were analyzed using the paired Wilcoxon test. Only statistically significant differences are indicated (*p < 0.05). (E, F) Cumulative probability plots show the percentage of induced LTD; negative values indicate LTP outcomes.

### Prenatal CBD exposure selectively compromises LTP in the male mPFC

To determine if prefrontal deficits extend to excitatory strengthening, we induced LTP using TBS, a canonical mechanism frequently disrupted in neuropsychiatric models (23,37–39). TBS reliably elicited LTP across all cohorts, elevating fEPSP amplitudes above baseline (Figure 8). In females, neither prenatal CBD nor THC altered LTP magnitude (Figure 8A, B, E). Conversely, a male-specific vulnerability emerged: prenatal CBD, but not THC, significantly blunted LTP amplitude compared to Sham controls (Figure 8C, D, F). This developmental profile reveals a sharp divergence between the two cannabinoids. While both ablate eCB-LTD, only CBD limits the capacity for LTP, specifically within the male lineage. This suggests that prenatal CBD drives a pervasive state of synaptic rigidity in males by impairing bidirectional plasticity (both strengthening and weakening), whereas the impact of THC is selectively restricted to the loss of synaptic depression.

**Figure 8:**
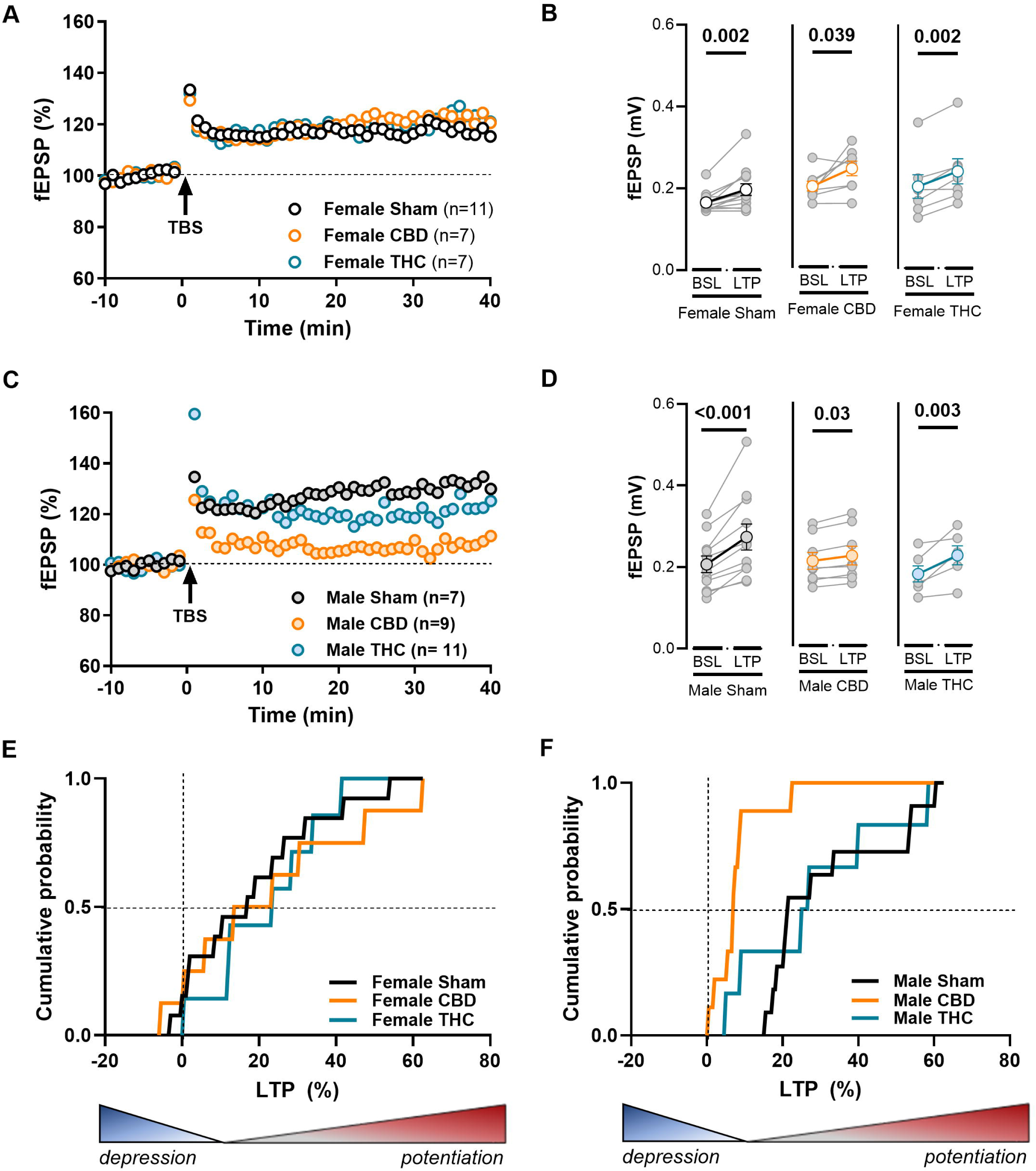
Prenatal cannabinoid exposure impairs long-term potentiation specifically in CBD-exposed males. (A, C) Representative time courses of normalized mean fEPSP amplitude before and after theta-burst stimulation (TBS). (B, D) Paired comparisons of mean absolute fEPSP amplitude during the 10-min baseline (BSL) versus the 30–40 min post-stimulation period (LTP). Individual experiments (grey dots) and group averages (colored circles) are shown. TBS reliably induces LTP across treatments in females (B) and in males exposed to Sham or THC (D). (E) Female offspring exposed to cannabinoids exhibit robust LTP with amplitude increases comparable to controls. (F) In contrast, CBD-exposed males show a marked reduction in LTP magnitude, with potentiation never exceeding ∼20%. Data are presented as mean ± SEM and were analyzed using the paired Wilcoxon test. Only statistically significant differences are indicated (*p < 0.05). (E, F) Cumulative probability plots illustrate the percentage of induced LTP.

### Sex-specific effects of gestational CBD on synaptic weighting and NMDAR Kinetics

Intact LTP following prenatal THC suggests functional preservation of NMDAR activation and AMPAR recruitment. Consequently, we isolated the cellular substrates underlying the male-specific synaptic rigidity unique to the CBD cohort. Basal synaptic efficacy and axonal excitability were unaltered, as input-output curves and maximal fEPSP amplitudes matched Sham controls in both sexes (Supplementary Fig. 5A-B). However, the postsynaptic AMPA/NMDA A/N) ratio was selectively elevated in CBD-exposed males (Figure 9A-B), indicating baseline functional saturation that occludes further potentiation. This postsynaptic occlusion coincided with disrupted NMDAR channel kinetics; gestational CBD significantly prolonged NMDA current rise times in males (Figure 9C), an effect poised to blunt the rapid calcium influx required to trigger LTP. Conversely, CBD females displayed accelerated NMDA decay kinetics (Figure 9D). Collectively, these findings demonstrate that male-specific LTP failure is driven by a dual synaptic deficit-functional saturation combined with a kinetic slowdown of NMDAR activation-that rigidly locks the prefrontal synapse and prevents further strengthening.

**Figure 9:**
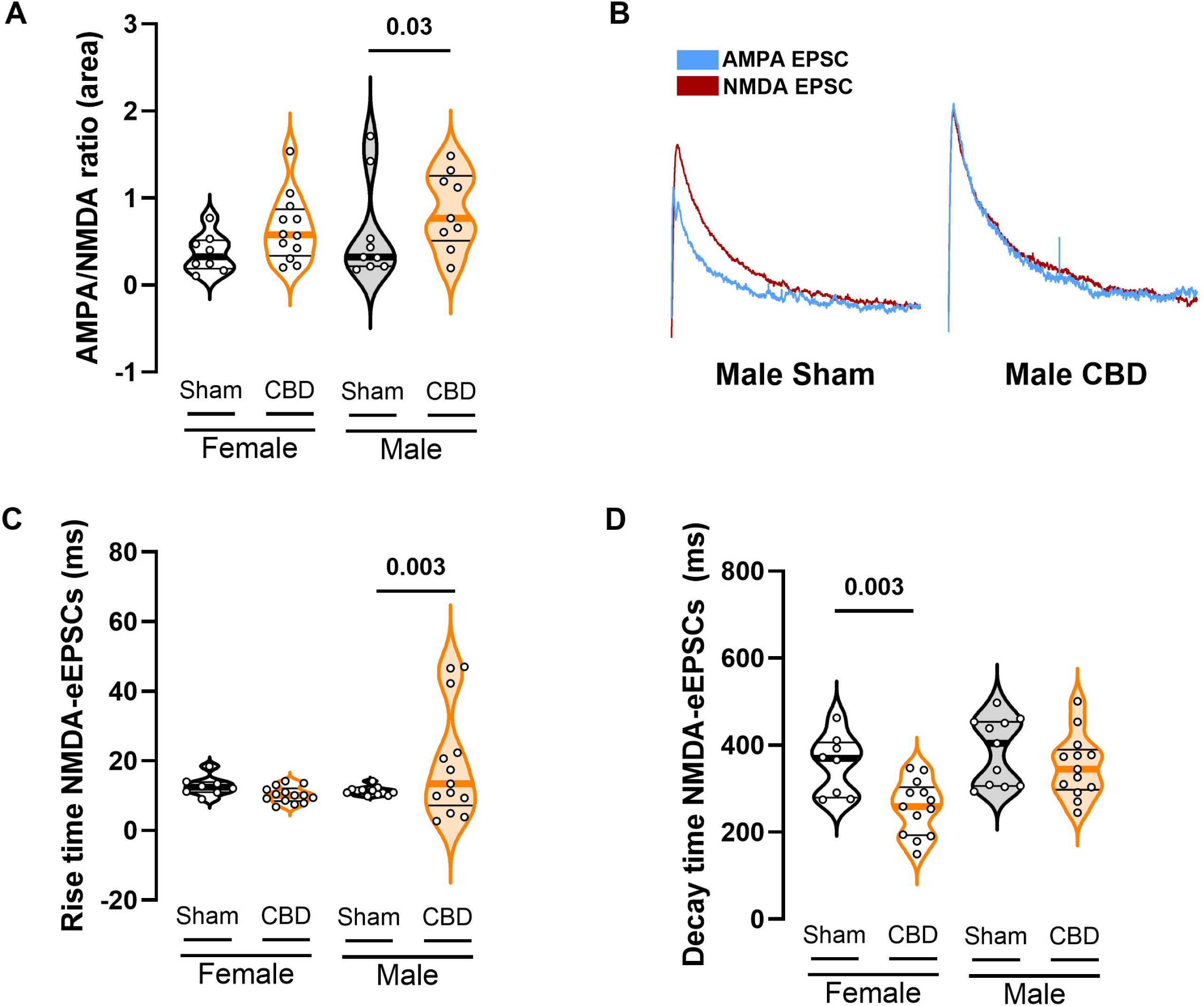
Selective increase in the AMPA/NMDA ratio in CBD-exposed male offspring. (A) CBD-exposed males show a significant increase in the AMPA/NMDA ratio in layer 5 pyramidal neurons of the mPFC. (B) Representative traces of NMDA-EPSCs (red) evoked at +30 mV in the presence of 50 µM NBQX, and AMPA-EPSCs (blue) obtained by digital subtraction of the NMDA component from the dual-component current at +30 mV. (C-D) CBD alters the kinetics of evoked NMDA currents. Male CBD-exposed offspring exhibit slower rise times, whereas CBD-exposed females display faster decay kinetics.. (A, C-D) Each point represents a single neuron; data are presented as violin plots (median and 25th–75th percentiles) and analyzed using two-way ANOVA followed by Šídák’s multiple-comparison test. Only statistically significant differences (p < 0.05) are indicated.

## Discussion

The data indicate that PCE alters prefrontal neurodevelopment, culminating in sex-specific rewiring in adulthood. Integrating ethological behavioral analysis with circuit-level electrophysiology shows that while THC and CBD converge on a universal loss of prefrontal eCB-LTD, they are associated with strikingly divergent synaptic signatures.

A central finding is the dissociation between classical anxiety and altered risk assessment. Neither drug altered EPM open-arm exploration (5,6); instead, changes were localized to the microstructure of risk assessment and behavioral switching. The concurrent presence of motor stereotypies (MB) and increased risk appraisal (SAP) points to a potential disruption in strategies required to resolve approach-avoidance conflicts (20–22,33). The female-specific SAP increase is associated with a hyper-vigilant state that may arise from a reduced prefrontal capacity to gate environmental information and resolve uncertainty (15,40), suggesting a targeted weakening of prefrontal behavioral strategy rather than a generalized affective shift.

Synaptically, both sexes and compounds converged on the complete ablation of eCB-LTD, a plausible cellular correlate to the universal increase in repetitive behaviors (MB). Because eCB-LTD is thought to participate in deselecting obsolete motor loops, its absence may contribute to a circuit-level environment where behavioral strategies appear less flexible. Consistent with other neurodevelopmental models (24,25,38,41,42), this loss of synaptic flexibility represents a shared endophenotype of PCE, indicating a lasting disruption in the maturation of endocannabinoid-dependent prefrontal plasticity (5,9,10).

The loss of eCB-LTD in THC-exposed female mice diverges from findings in rats, where this plasticity was preserved in females (5). This discrepancy may reflect species-specific ontogenetic windows, where a more concentrated prefrontal eCB maturation period (GD5–18) potentially increases mouse vulnerability. Alternatively, divergent endocannabinoid mediated plasticity(43) profiles might suggest the female mouse lacks homeostatic buffers that could otherwise protect receptor coupling.

While eCB-LTD loss appeared universal, plasticity patterns diverged sharply in males. Prenatal THC did not affect LTP, suggesting intact excitatory strengthening. Conversely, male CBD exposure was uniquely associated with reduced synaptic flexibility, characterized by a bidirectional limitation where neither eCB-LTD nor NMDA-LTP was observed. Mechanistically, this state may be linked to elevated AMPA/NMDA ratios and slowed NMDAR activation kinetics, changes which could represent functional synaptic saturation. This ceiling effect may constrain prefrontal function. Since L5 pyramidal neurons are the primary mPFC output, this rigidity suggests final executive commands may be less adaptable before reaching downstream targets like the striatum or amygdala. This constraint appears to limit the plasticity typically associated with flexibility, suggesting a distinct male sensitivity to CBD.

Reconciling male synaptic pathology with behavior, CBD-exposed males lacked the female-specific SAP phenotype, showing only the universal MB increase. This dissociation may reflect the nature of the male deficit: a bidirectional plasticity collapse and elevated AMPA/NMDA ratios suggest functional saturation rather than female dynamic instability. While eCB-LTD loss predicts perseverative MB loops, this locked state may preclude nuanced risk-assessment strategies requiring flexible arbitration. Additionally, control males inherently default to neutral postures, potentially masking treatment-induced shifts. The absence of a unique male SAP phenotype thus does not contradict rigidity, but points to a ceiling effect restricting adaptation to a loss of flexibility (MB) rather than a strategic shift (SAP).

Conversely, female offspring exhibited divergent reorganizations of E/I balance and throughput. CBD-exposed females displayed a homeostatic scaled-up architecture with coordinated amplification of both excitatory and inhibitory drives. While preserving net E/I balance, this scaling is associated with elevated throughput that may correlate with unique SAP deficits, where altered circuit dynamics could contribute to the hyper-vigilant risk-appraisal strategy observed behaviorally. In contrast, female THC exposure was associated with a pro-excitatory shift, reflecting circuit disinhibition. This divergence suggests that the mPFC undergoes a selective reorganization of intrinsic excitability and synaptic integration depending critically on compound identity.

In CBD-exposed females, the scaled-up architecture maintains a balanced E/I ratio. While this compensation preserves equilibrium, it may alter the dynamics required for flexibility. The circuit is paradoxically balanced yet functionally rigid, manifesting as the observed hyper-vigilant behaviors. We propose that this homeostatic stabilization occurs at the expense of the circuit’s ability to adapt to the outside world.

These findings challenge the perception of CBD as a benign alternative to THC. Given rising gestational consumption, our data carry important translational considerations, suggesting CBD is associated with discrete, sex-specific molecular signatures, specifically, male synaptic rigidity and altered female circuit dynamics. This specificity underscores the value of sex-disaggregated analyses in cannabinoid research. In sum, prenatal cannabinoid exposure appears to modify the fundamental rules of synaptic integration and plasticity rather than merely dampening or exciting cortical activity. By mapping universal eCB-LTD loss against sex-specific trajectories, this study offers a biological framework for considering the long-term risks of gestational cannabinoid use.

## Author Contributions

A.C.-R.: conceptualization, data curation, formal analysis, validation, writing (review and editing). D.I.: data curation, writing (review and editing). O.L.: D: data curation, formal analysis. S.W: formal analysis; A.D.: methodology. P.C.: conceptualization, methodology, project administration, supervision. O.J.J.M.: conceptualization, supervision, funding acquisition, methodology, project administration, writing (original draft, review, and editing). All authors have read and agreed to the published version of the manuscript.

## Acknowledgements

The authors are grateful to the Chavis-Manzoni team members for helpful discussions. The authors would like to acknowledge the use of the large language model Gemini (Google) for structural and linguistic refinements. Anna E. Dudek and Shuai Wang acknowledge the financial support provided by the French government under the France 2030 investment plan, through the Initiative d’Excellence d’Aix-Marseille Université – AMIDEX (grant no. AMX-22-CEX-057).

## Supplementary figures legends

**Supplementary Figure 1: Prenatal cannabinoid (THC or CBD) exposure does not affect open-arm time in the Elevated Plus Maze in adult offspring.** (A) Heatmap showing the time spent in each arm of the maze over a 300-second trial. Warm colors (e.g., red) indicate areas visited most frequently, while cool colors (e.g., blue) denote areas visited least frequently. Quantitative analyses include (B) total distance moved, (C) preference for open arms, (D) time spent in the closed arm and center, and (E) time spent in the open arms. (B) Notably, THC-exposed female offspring displayed increased locomotor activity, as reflected in increased distance moved. (C-E) However, analysis of relative time spent in different maze zones revealed no significant differences between groups, regardless of sex or prenatal exposure to CBD or THC violin plots (median and 25th–75th percentiles) and analyzed using two-way ANOVA followed by Šídák’s multiple-comparison test. No significant effects were detected. Group sizes were as follows Sham female (N = 13), CBD female (N = 12), THC female (N = 16), Sham male (N = 11), CBD male (N = 10), THC male (N = 16).

**Supplementary Figure 2: Prenatal cannabinoid exposure induces sex-specific changes in the passive properties of mPFC principal neurons.** (A) Representative firing patterns evoked by hyperpolarizing and depolarizing current injections (500 ms; −400 to +150 pA in 50-pA steps) in Sham-, CBD-, and THC-exposed offspring of both sexes. Recordings were obtained from layer 5 pyramidal neurons of the mPFC (Bregma +2.34). (B-D) Quantitative analysis of passive membrane properties of mPFC pyramidal neurons. (B) Resting membrane potential was not altered. (C) Progressive depolarizing current injections (10-pA steps) reveal an increased rheobase in females prenatally exposed to THC. (E) Capacitance, an indirect measure of cell size, remain similar across conditions. (E-F) No significant differences were observed in current–voltage (I–V) relationships of mPFC pyramidal neurons across sex or treatment. Responses were elicited by incremental current injections ranging from −400 to +200 pA in 50-pA steps. (B–D) Each dot represents a single neuron; data are presented as violin plots (median, and 25th–75th percentiles) and were analyzed using two-way ANOVA followed by Šídák’s multiple-comparison test. (E-F) Each dot represents the average response of all neurons at each current step; data are shown as mean ±SEM and analyzed using multiple Mann–Whitney U tests. Statistical significance (p-value < 0.05) is indicated in the graphs and in Table 6. Sample sizes (# neurons / # animals) were females Sham (34/9), CBD (31/10), THC (26/9); males Sham (27/8), CBD (34/14), THC (30/8).

**Supplementary Figure 3: Prenatal cannabinoid exposure alters excitatory synaptic transmission in female progeny.** (A) On average, CBD-exposed females exhibit larger sEPSC amplitudes, while (B) sEPSC frequency is increased in CBD-exposed females compared with Sham and THC groups. (A, B) Data are presented as violin plots (median, and 25th–75th percentiles) and were analyzed using two-way ANOVA followed by Šídák’s correction for multiple comparisons. Only statistically significant differences (p < 0.05) are indicated in the graphs additional statistical information can be found in Table 9. Sample sizes (# neurons / # animals) were females Sham (24/9), CBD (26/12), THC (21/9); males Sham (21/8), CBD (21/13), THC (27/8).

**Supplementary Figure 4: Prenatal cannabinoid exposure differentially alters inhibitory synaptic transmission in female and male progeny.** (A) On average, CBD-exposed females exhibit larger sIPSC amplitudes, while prenatal THC decreases the mean inhibitory transmission in males. Despite the changes on the current amplitude, (B) Mean sIPSC frequency does not change in cannabinoid-treated females. (A, B) Data are presented as violin plots (median, and 25th–75th percentiles) and were analyzed using two-way ANOVA followed by Šídák’s correction for multiple comparisons. (D, E, G, H) The graphs display the relative distribution of all events in 6min recording, these data were analyzed using log-normal curve fitting with confidence intervals (± CI). Only statistically significant differences (p < 0.05) are indicated in the graphs. Further statistical details can be found on Table 6. Sample sizes (# neurons / # animals) were females Sham (20/6), CBD (27/7), THC (21/5); males Sham (27/7), CBD (21/7), THC (24/5).

**Supplementary Figure 5: Averaged fEPSP amplitudes plotted as a function of stimulus intensity.** (A,B) Synaptic strength remains stable across treatments, indicating that baseline recruitment does not account for the observed plasticity differences. Each dot represents the average response of all neurons at each current step; data are shown as mean ±SEM and analyzed using multiple Mann–Whitney U tests.

## References

1. Bhatia D, Battula S, Mikulich-Gilbertson S, Sakai J, Hammond D. Cannabidiol-Only Product Use in Pregnancy in the United States and Canada: Findings From the International Cannabis Policy Study. Obstetrics & Gynecology. 2024 Aug;144(2):156–9. doi:10.1097/AOG.0000000000005603

2. Bara A, Ferland JMN, Rompala G, Szutorisz H, Hurd YL. Cannabis and synaptic reprogramming of the developing brain. Nat Rev Neurosci. 2021 Jul;22(7):423–38. doi:10.1038/s41583-021-00465-5

3. Harkany T, Guzmán M, Galve-Roperh I, Berghuis P, Devi LA, Mackie K. The emerging functions of endocannabinoid signaling during CNS development. Trends Pharmacol Sci. 2007 Feb;28(2):83–92. doi:10.1016/j.tips.2006.12.004 PubMed PMID: 17222464.

4. Harkany T, Cinquina V. Physiological Rules of Endocannabinoid Action During Fetal and Neonatal Brain Development. Cannabis Cannabinoid Res. 2021 Oct;6(5):381–8. doi:10.1089/can.2021.0096 PubMed PMID: 34619043; PubMed Central PMCID: PMC8664114.

5. Bara A, Manduca A, Bernabeu A, Borsoi M, Serviado M, Lassalle O, et al. Sex-dependent effects of in utero cannabinoid exposure on cortical function. Elife. 2018 Sep 11;7:e36234. doi:10.7554/eLife.36234 PubMed PMID: 30201092; PubMed Central PMCID: PMC6162091.

6. Manduca A, Servadio M, Melancia F, Schiavi S, Manzoni OJ, Trezza V. Sex-specific behavioural deficits induced at early life by prenatal exposure to the cannabinoid receptor agonist WIN55, 212-2 depend on mGlu5 receptor signalling. Br J Pharmacol. 2020 Jan;177(2):449–63. doi:10.1111/bph.14879 PubMed PMID: 31658362; PubMed Central PMCID: PMC6989958.

7. Vargish GA, Pelkey KA, Yuan X, Chittajallu R, Collins D, Fang C, et al. Persistent inhibitory circuit defects and disrupted social behaviour following in utero exogenous cannabinoid exposure. Mol Psychiatry. 2017 Jan;22(1):56–67. doi:10.1038/mp.2016.17 PubMed PMID: 26976041; PubMed Central PMCID: PMC5025333.

8. Frau R, Miczán V, Traccis F, Aroni S, Pongor CI, Saba P, et al. Prenatal THC exposure produces a hyperdopaminergic phenotype rescued by pregnenolone. Nat Neurosci. 2019 Dec;22(12):1975–85. doi:10.1038/s41593-019-0512-2

9. Scheyer AF, Borsoi M, Pelissier-Alicot AL, Manzoni OJJ. Maternal Exposure to the Cannabinoid Agonist WIN 55,12,2 during Lactation Induces Lasting Behavioral and Synaptic Alterations in the Rat Adult Offspring of Both Sexes. eNeuro. 2020;7(5):ENEURO.0144-20.2020. doi:10.1523/ENEURO.0144-20.2020 PubMed PMID: 32868310; PubMed Central PMCID: PMC7540927.

10. Scheyer AF, Borsoi M, Pelissier-Alicot AL, Manzoni OJJ. Perinatal THC exposure via lactation induces lasting alterations to social behavior and prefrontal cortex function in rats at adulthood. Neuropsychopharmacol. 2020 Oct;45(11):1826–33. doi:10.1038/s41386-020-0716-x

11. Swenson KS, Gomez Wulschner LE, Hoelscher VM, Folts L, Korth KM, Oh WC, et al. Fetal cannabidiol (CBD) exposure alters thermal pain sensitivity, problem-solving, and prefrontal cortex excitability. Mol Psychiatry. 2023 Aug;28(8):3397–413. doi:10.1038/s41380-023-02130-y

12. Iezzi D, Caceres-Rodriguez A, Chavis P, Manzoni OJJ. In utero exposure to cannabidiol disrupts select early-life behaviors in a sex-specific manner. Transl Psychiatry. 2022 Dec 5;12(1):501. doi:10.1038/s41398-022-02271-8

13. Maciel IDS, Abreu GHDD, Johnson CT, Bonday R, Bradshaw HB, Mackie K, et al. Perinatal CBD or THC Exposure Results in Lasting Resistance to Fluoxetine in the Forced Swim Test: Reversal by Fatty Acid Amide Hydrolase Inhibition. Cannabis and Cannabinoid Research. 2022 Jun 1;7(3):318–27. doi:10.1089/can.2021.0015

14. Iezzi D, Cáceres-Rodríguez A, Pereira-Silva J, Chavis P, Manzoni OJJ. Gestational CBD Shapes Insular Cortex in Adulthood. Cells. 2024 Sep 4;13(17):1486. doi:10.3390/cells13171486 PubMed PMID: 39273056; PubMed Central PMCID: PMC11394289.

15. Iezzi D, Cáceres-Rodríguez A, Chavis P, Manzoni OJ. Sex-specific disruptions in the developmental trajectory of anxiety-like behaviors due to prenatal cannabidiol exposure. Transl Psychiatry. 2025 Oct 6;15(1):354. doi:10.1038/s41398-025-03517-x PubMed PMID: 41053015; PubMed Central PMCID: PMC12500939.

16. Halladay LR, Blair HT. Distinct ensembles of medial prefrontal cortex neurons are activated by threatening stimuli that elicit excitation vs. inhibition of movement. J Neurophysiol. 2015 Aug;114(2):793–807. doi:10.1152/jn.00656.2014 PubMed PMID: 25972588; PubMed Central PMCID: PMC4533059.

17. Albrechet-Souza L, Borelli KG, Brandão ML. Activity of the medial prefrontal cortex and amygdala underlies one-trial tolerance of rats in the elevated plus-maze. J Neurosci Methods. 2008 Mar 30;169(1):109–18. doi:10.1016/j.jneumeth.2007.11.025 PubMed PMID: 18190969.

18. Klune CB, Goodpaster CM, Gongwer MW, Gabriel CJ, An J, Chen R, et al. Developmentally distinct architectures in top-down pathways controlling threat avoidance. Nat Neurosci. 2025 Apr;28(4):823–35. doi:10.1038/s41593-025-01890-w PubMed PMID: 39972221; PubMed Central PMCID: PMC11978489.

19. de Brouwer G, Fick A, Harvey BH, Wolmarans DW. A critical inquiry into marble-burying as a preclinical screening paradigm of relevance for anxiety and obsessive-compulsive disorder: Mapping the way forward. Cogn Affect Behav Neurosci. 2019 Feb;19(1):1–39. doi:10.3758/s13415-018-00653-4 PubMed PMID: 30361863.

20. Rodgers RJ, Dalvi A. Anxiety, defence and the elevated plus-maze. Neurosci Biobehav Rev. 1997 Nov;21(6):801–10. doi:10.1016/s0149-7634(96)00058-9 PubMed PMID: 9415905.

21. Walf AA, Frye CA. The use of the elevated plus maze as an assay of anxiety-related behavior in rodents. Nat Protoc. 2007;2(2):322–8. doi:10.1038/nprot.2007.44 PubMed PMID: 17406592; PubMed Central PMCID: PMC3623971.

22. Thomas A, Burant A, Bui N, Graham D, Yuva-Paylor LA, Paylor R. Marble burying reflects a repetitive and perseverative behavior more than novelty-induced anxiety. Psychopharmacology (Berl). 2009 Jun;204(2):361–73. doi:10.1007/s00213-009-1466-y PubMed PMID: 19189082; PubMed Central PMCID: PMC2899706.

23. Thomazeau A, Lassalle O, Iafrati J, Souchet B, Guedj F, Janel N, et al. Prefrontal deficits in a murine model overexpressing the down syndrome candidate gene dyrk1a. J Neurosci. 2014 Jan 22;34(4):1138–47. doi:10.1523/JNEUROSCI.2852-13.2014 PubMed PMID: 24453307; PubMed Central PMCID: PMC3953590.

24. Lafourcade M, Larrieu T, Mato S, Duffaud A, Sepers M, Matias I, et al. Nutritional omega-3 deficiency abolishes endocannabinoid-mediated neuronal functions. Nat Neurosci. 2011 Mar;14(3):345–50. doi:10.1038/nn.2736 PubMed PMID: 21278728.

25. Jung KM, Sepers M, Henstridge CM, Lassalle O, Neuhofer D, Martin H, et al. Uncoupling of the endocannabinoid signalling complex in a mouse model of fragile X syndrome. Nat Commun. 2012;3:1080. doi:10.1038/ncomms2045 PubMed PMID: 23011134; PubMed Central PMCID: PMC3657999.

26. Bouamrane L, Scheyer AF, Lassalle O, Iafrati J, Thomazeau A, Chavis P. Reelin-Haploinsufficiency Disrupts the Developmental Trajectory of the E/I Balance in the Prefrontal Cortex. Front Cell Neurosci. 2016;10:308. doi:10.3389/fncel.2016.00308 PubMed PMID: 28127276; PubMed Central PMCID: PMC5226963.

27. Iafrati J, Malvache A, Gonzalez Campo C, Orejarena MJ, Lassalle O, Bouamrane L, et al. Multivariate synaptic and behavioral profiling reveals new developmental endophenotypes in the prefrontal cortex. Sci Rep. 2016 Oct 21;6:35504. doi:10.1038/srep35504 PubMed PMID: 27765946; PubMed Central PMCID: PMC5073243.

28. Labouesse MA, Lassalle O, Richetto J, Iafrati J, Weber-Stadlbauer U, Notter T, et al. Hypervulnerability of the adolescent prefrontal cortex to nutritional stress via reelin deficiency. Mol Psychiatry. 2017 Jul;22(7):961–71. doi:10.1038/mp.2016.193 PubMed PMID: 27843148.

29. Bolhuis K, Kushner SA, Yalniz S, Hillegers MHJ, Jaddoe VWV, Tiemeier H, et al. Maternal and paternal cannabis use during pregnancy and the risk of psychotic-like experiences in the offspring. Schizophr Res. 2018 Dec;202:322–7. doi:10.1016/j.schres.2018.06.067 PubMed PMID: 29983267.

30. El Marroun H, Brown QL, Lund IO, Coleman-Cowger VH, Loree AM, Chawla D, et al. An epidemiological, developmental and clinical overview of cannabis use during pregnancy. Prev Med. 2018 Nov;116:1–5. doi:10.1016/j.ypmed.2018.08.036 PubMed PMID: 30171964.

31. Grant KS, Petroff R, Isoherranen N, Stella N, Burbacher TM. Cannabis use during pregnancy: Pharmacokinetics and effects on child development. Pharmacol Ther. 2018 Feb;182:133–51. doi:10.1016/j.pharmthera.2017.08.014 PubMed PMID: 28847562; PubMed Central PMCID: PMC6211194.

32. Sarikahya MH, Cousineau SL, De Felice M, Szkudlarek HJ, Wong KKW, DeVuono MV, et al. Prenatal THC exposure induces long-term, sex-dependent cognitive dysfunction associated with lipidomic and neuronal pathology in the prefrontal cortex-hippocampal network. Mol Psychiatry. 2023 Oct;28(10):4234–50. doi:10.1038/s41380-023-02190-0 PubMed PMID: 37525013.

33. Blanchard DC, Griebel G, Pobbe R, Blanchard RJ. Risk assessment as an evolved threat detection and analysis process. Neurosci Biobehav Rev. 2011 Mar;35(4):991–8. doi:10.1016/j.neubiorev.2010.10.016 PubMed PMID: 21056591.

34. Xu P, Chen A, Li Y, Xing X, Lu H. Medial prefrontal cortex in neurological diseases. Physiol Genomics. 2019 Sep 1;51(9):432–42. doi:10.1152/physiolgenomics.00006.2019 PubMed PMID: 31373533; PubMed Central PMCID: PMC6766703.

35. Dembrow NC, Zemelman BV, Johnston D. Temporal dynamics of L5 dendrites in medial prefrontal cortex regulate integration versus coincidence detection of afferent inputs. J Neurosci. 2015 Mar 18;35(11):4501–14. doi:10.1523/JNEUROSCI.4673-14.2015 PubMed PMID: 25788669; PubMed Central PMCID: PMC4363381.

36. Lee AT, Gee SM, Vogt D, Patel T, Rubenstein JL, Sohal VS. Pyramidal neurons in prefrontal cortex receive subtype-specific forms of excitation and inhibition. Neuron. 2014 Jan 8;81(1):61–8. doi:10.1016/j.neuron.2013.10.031 PubMed PMID: 24361076; PubMed Central PMCID: PMC3947199.

37. Martin HGS, Lassalle O, Brown JT, Manzoni OJ. Age-Dependent Long-Term Potentiation Deficits in the Prefrontal Cortex of the Fmr1 Knockout Mouse Model of Fragile X Syndrome. Cereb Cortex. 2016 May;26(5):2084–92. doi:10.1093/cercor/bhv031 PubMed PMID: 25750254.

38. Manduca A, Bara A, Larrieu T, Lassalle O, Joffre C, Layé S, et al. Amplification of mGlu5-Endocannabinoid Signaling Rescues Behavioral and Synaptic Deficits in a Mouse Model of Adolescent and Adult Dietary Polyunsaturated Fatty Acid Imbalance. J Neurosci. 2017 Jul 19;37(29):6851–68. doi:10.1523/JNEUROSCI.3516-16.2017 PubMed PMID: 28630250; PubMed Central PMCID: PMC6705718.

39. Kauer JA, Malenka RC. Synaptic plasticity and addiction. Nat Rev Neurosci. 2007 Nov;8(11):844–58. doi:10.1038/nrn2234 PubMed PMID: 17948030.

40. Holly KS, Orndorff CO, Murray TA. MATSAP: An automated analysis of stretch-attend posture in rodent behavioral experiments. Sci Rep. 2016 Aug 9;6:31286. doi:10.1038/srep31286 PubMed PMID: 27503239; PubMed Central PMCID: PMC4977506.

41. Bosch-Bouju C, Larrieu T, Linders L, Manzoni OJ, Layé S. Endocannabinoid-Mediated Plasticity in Nucleus Accumbens Controls Vulnerability to Anxiety after Social Defeat Stress. Cell Rep. 2016 Aug 2;16(5):1237–42. doi:10.1016/j.celrep.2016.06.082 PubMed PMID: 27452462.

42. Martin HGS, Lassalle O, Manzoni OJ. Differential Adulthood Onset mGlu5 Signaling Saves Prefrontal Function in the Fragile X Mouse. Cereb Cortex. 2017 Dec 1;27(12):5592–602. doi:10.1093/cercor/bhw328 PubMed PMID: 27797833.

43. Bernabeu A, Bara A, Murphy Green MN, Manduca A, Wager-Miller J, Borsoi M, et al. Sexually Dimorphic Adolescent Trajectories of Prefrontal Endocannabinoid Synaptic Plasticity Equalize in Adulthood, Reflected by Endocannabinoid System Gene Expression. Cannabis & Cannabinoid Research. 2023 Oct 1;8(5):749–67. doi:10.1089/can.2022.0308

