## Supplementary figures and images for "Prenatal Cannabidiol and Δ9-Tetrahydrocannabinol exposure lead to sex-specific disruptions in risk assessment and behavioral switching via divergent rewiring of the adult mPFC"

### sup1

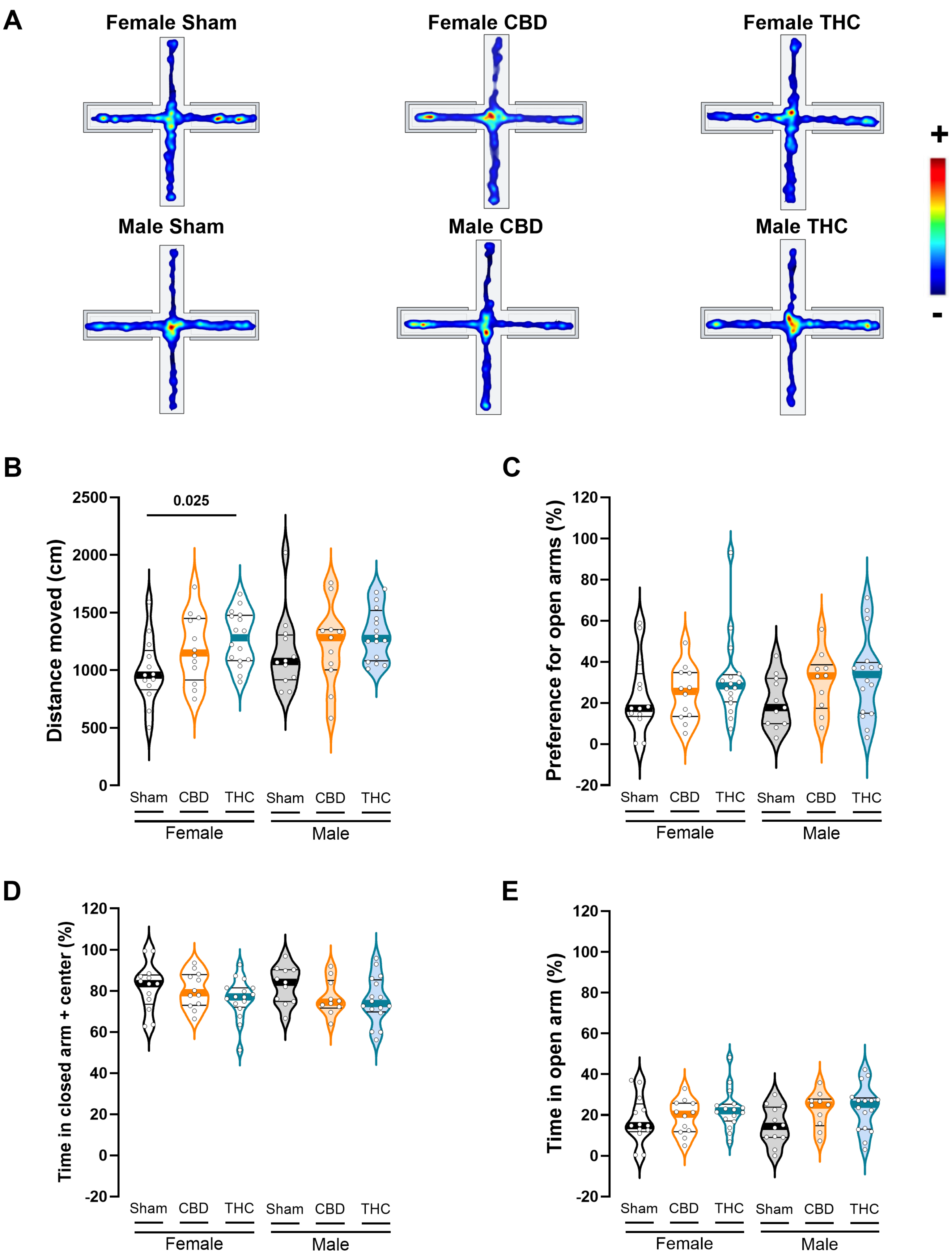

### sup2

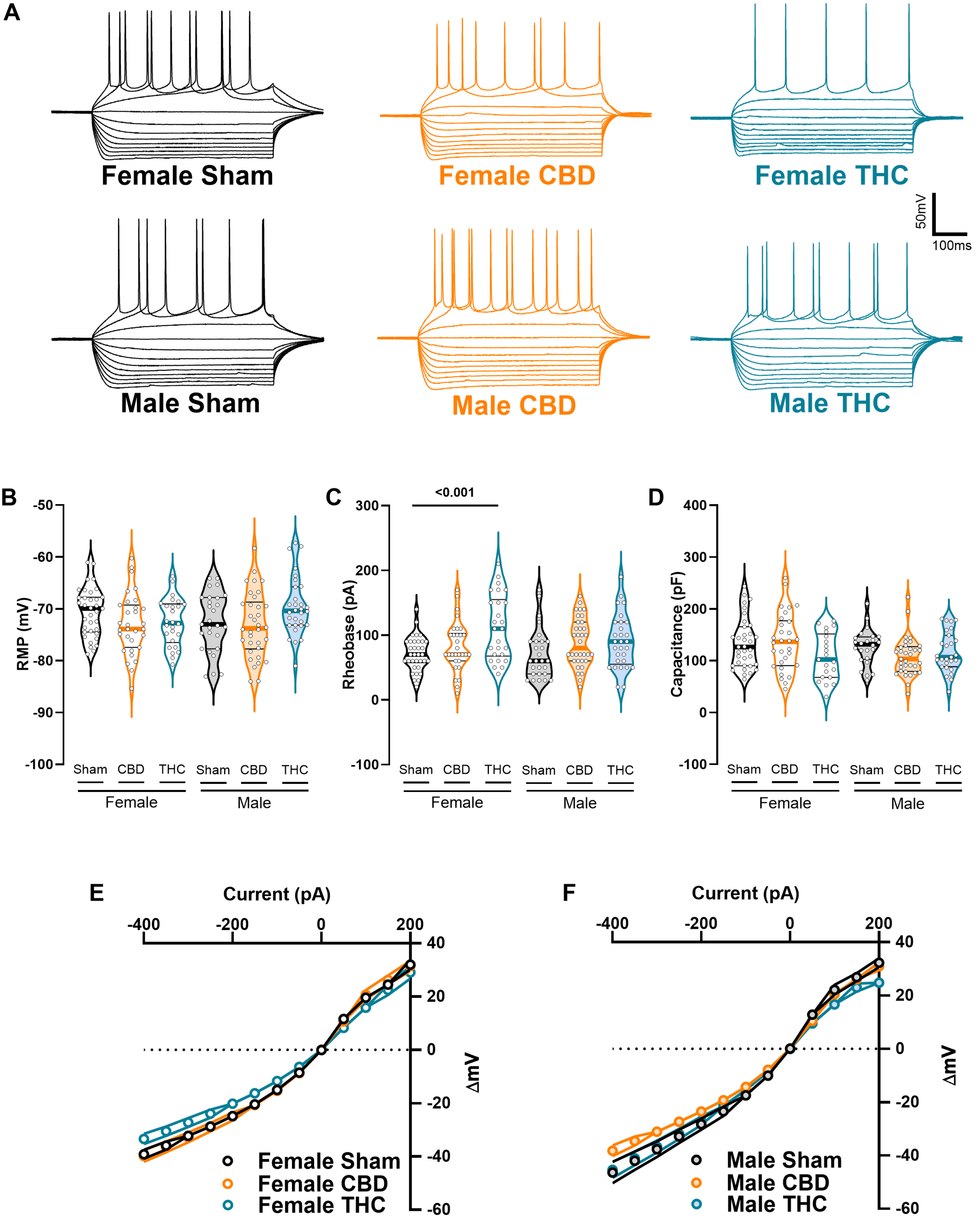

### sup3

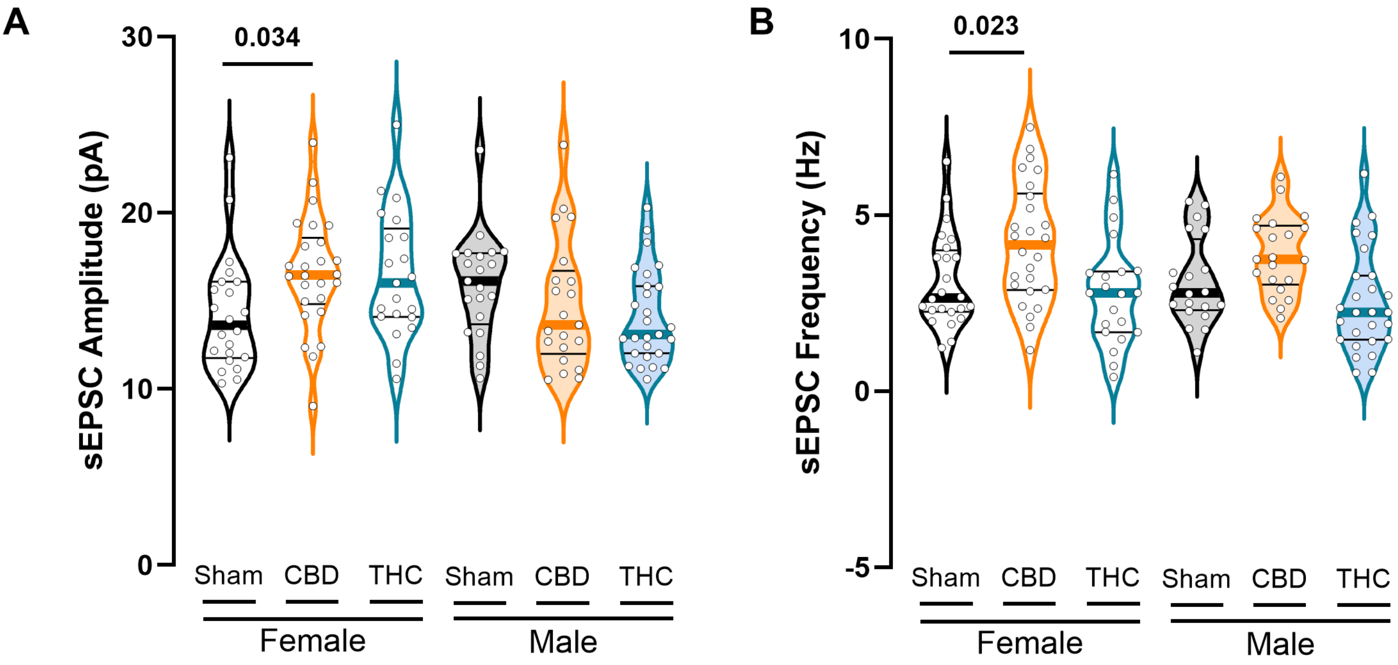

### sup4

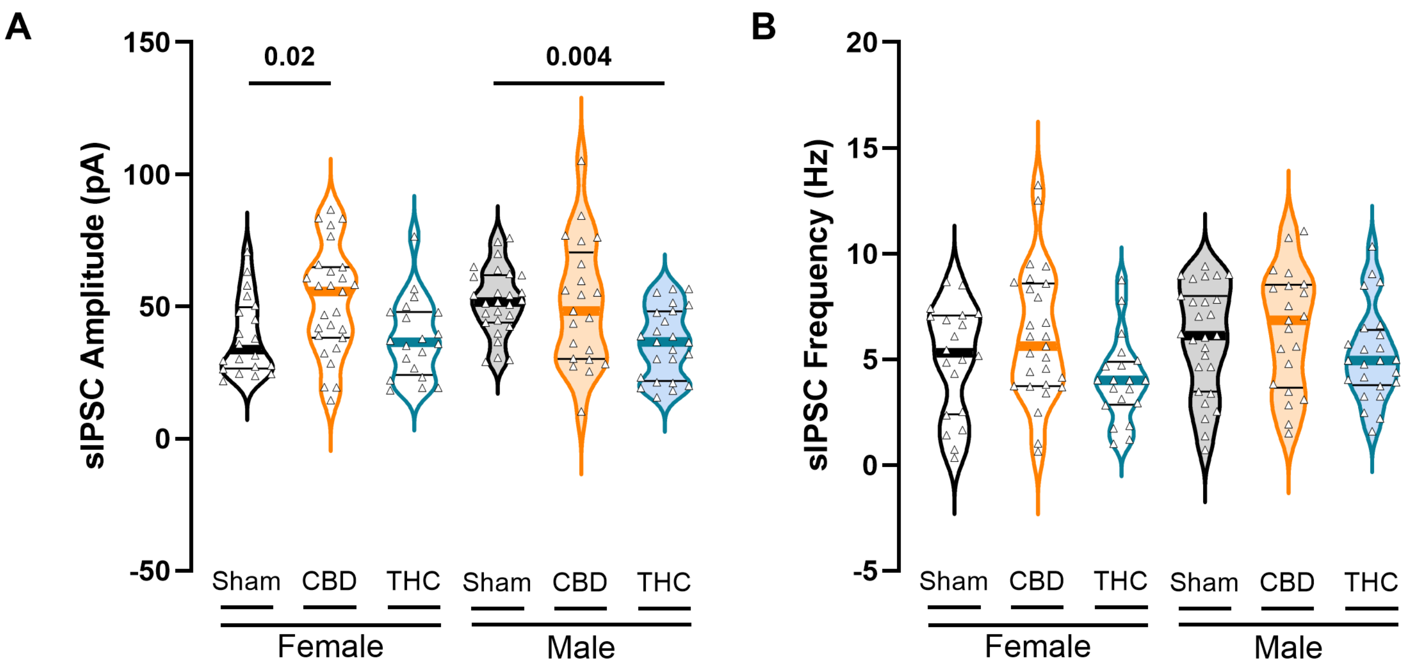

### sup5

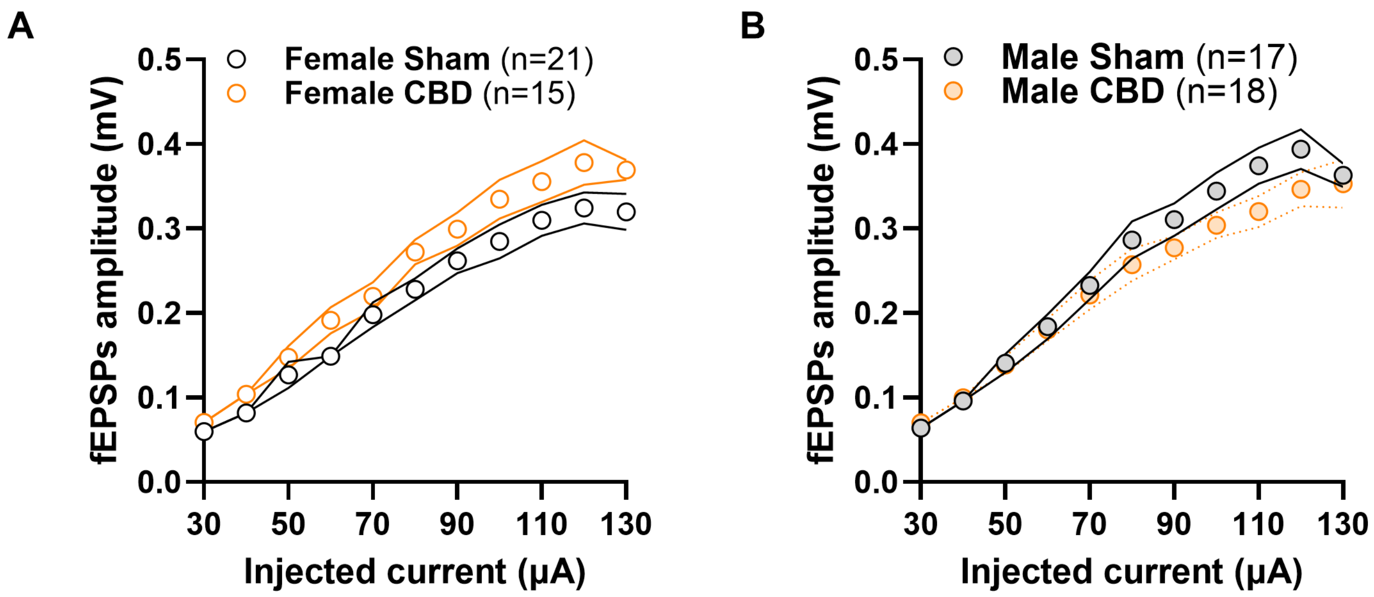
